# Anthropogenic disturbance drives dispersal syndromes, demography, and gene flow in spatially structured amphibian populations

**DOI:** 10.1101/789966

**Authors:** Hugo Cayuela, Aurélien Besnard, Julien Cote, Martin Laporte, Eric Bonnaire, Julian Pichenot, Nicolas Schtickzelle, Arnaud Bellec, Pierre Joly, Jean-Paul Léna

## Abstract

There is growing evidence that anthropogenic landscapes can strongly influence the evolution of dispersal, particularly through fragmentation, and may drive organisms into an evolutionary trap by suppressing dispersal. However, the influence on dispersal evolution of anthropogenic variation in habitat patch turnover has so far been largely overlooked. In this study, we examined how human-driven variation in patch persistence affects dispersal rates and distances, determines dispersal-related phenotypic specialization, and drives neutral genetic structure in spatially structured populations. We addressed this issue in an amphibian, *Bombina variegata*, using an integrative approach combining capture–recapture modeling, demographic simulation, common garden experiments, and population genetics. *B. variegata* reproduces in small ponds that occur either in habitat patches that are persistent (i.e. several decades or more), located in riverine environments with negligible human activity, or in patches that are highly temporary (i.e. a few years), created by logging operations in intensively harvested woodland. Our capture–recapture models revealed that natal and breeding dispersal rates and distances were drastically higher in spatially structured populations (SSPs) in logging environments than in riverine SSPs. Population simulations additionally showed that dispersal costs and benefits drive the fate of logging SSPs, which cannot persist without dispersal. The common garden experiments revealed that toadlets reared in laboratory conditions have morphological and behavioral specialization that depends on their habitat of origin. Toadlets from logging SSPs were found to have higher boldness and exploration propensity than those from riverine SSPs, indicating transgenerationally transmitted dispersal syndromes. We also found contrasting patterns of neutral genetic diversity and gene flow in riverine and logging SSPs, with genetic diversity and effective population size considerably higher in logging than in riverine SSPs. In parallel, intra-patch inbreeding and relatedness levels were lower in logging SSPs. Controlling for the effect of genetic drift and landscape connectivity, gene flow was found to be higher in logging than in riverine SSPs. Taken together, these results indicate that anthropogenic variation in habitat patch turnover may have an effect at least as important as landscape fragmentation on dispersal evolution and the long-term viability and genetic structure of wild populations.

## Introduction

In the age of the Anthropocene, a significant proportion of land cover has been replaced by human-dominated landscapes (Foley et al. 2005, Pereira et al. 2010, Gibson et al. 2011, Tilman et al. 2017), with the result that the conditions prevailing in these anthropogenic environments now shape the evolutionary course of almost all species (Otto 2018, Pelletier & Coltman 2018). Land use changes usually have the simultaneous effects of habitat loss, alteration, and/or fragmentation into small habitat patches isolated in a more or less hostile matrix (i.e. unsuitable habitat) (Fahrig 2003, Villard & Metzger 2014). While it is widely accepted that habitat loss is the main factor involved in local extinction and biodiversity loss (Sala et al. 2000, Pereira et al. 2010, Newbold et al. 2015, Tilman et al. 2017), it is also increasingly recognized that habitat alteration and fragmentation are critical ecological and evolutionary drivers in anthropogenic landscapes (Villard & Metzger 2014, Haddad et al 2015). Interestingly, the risk of extirpation from anthropogenic landscapes appears to differ between species (Frishkoff et al. 2014, Edward et al. 2015, Nowakoski et al. 2018) and research is needed to identify the phenotypic traits that allow some species to cope with and succeed in human-dominated contexts.

Dispersal, i.e. the movement from birth to breeding patch (natal dispersal) or between successive breeding patches (breeding dispersal), is a key ecological and evolutionary process. Dispersal provides the demographic supply for population rescuing, habitat (re)colonization (Hanski & Gaggiotti 2004, Bowler & Benton 2005, Gilpin 2012), and range expansion (Travis et al. 2009, Kubisch et al. 2014, Ochocki & Miller 2017). Furthermore, it determines the intensity and direction of gene flow, which has far-reaching consequences for local genetic diversity and adaptive processes (Lenormand 2002, Ronce 2007, Petit & Broquet 2009, Cayuela et al. 2018a). Dispersal is a complex phenotype, partially controlled by genetics and relying on a suite of morphological, behavioral and life history traits that may be subject to joint selection (Saastamoinen et al. 2018). Such associations between dispersal and individual phenotype are called ‘dispersal syndromes’ and lead to phenotypic specialization within and between populations (Matthysen 2012, Cote et al. 2010, Ronce & Clobert 2012). Dispersal is also influenced by patch and landscape characteristics: individuals are expected to adjust their dispersal decisions according to the fitness prospects of a patch (i.e. ‘informed dispersal’, Clobert et al. 2009), leading to context-dependent dispersal.

Over the last two decades, an increasing number of studies have suggested that landscape anthropization is an important determinant in dispersal evolution as it affects the balance between fitness benefits and the direct and indirect costs of moving (Bonte et al. 2012) incurred at the different stages of the dispersal process (i.e. emigration, transience, and immigration) (Kokko & Lopez-Sepulcre 2006, Cheptou et al. 2017, Cote et al. 2017). The majority of efforts have been devoted to better understanding the influence of habitat fragmentation on dispersal evolution in anthropogenic landscapes (Cheptou et al. 2017, Cote et al. 2017, Legrand et al. 2017, Atkin et al. 2019). Overall, these studies have reported reduced dispersal propensity or capacity in fragmented landscapes, which is usually attributed to prohibitive costs during the transition phase across the matrix (Cheptou et al. 2017, Cote et al 2017). This hypothesis is supported by landscape genetic studies, which often report increased spatial genetic differentiation depending on the harshness of the matrix that separates demes (Baguette et al. 2013, Cushman et al. 2015). Nevertheless, habitat fragmentation may have a contradictory effect on the evolution of dispersal: on one hand, it may make transition across the matrix costlier, but on the other, it may make dispersal profitable due to the increased local extinction risk caused by heightened demographic stochasticity within severely fragmented landscapes (Ronce 2007, Hanski & Mononen 2011, Cote et al. 2017). The best empirical support of this hypothesis is perhaps the well-documented selection for dispersal-specialized phenotypes observed in spatially structured populations (SSPs, Thomas & Kunin 1999) of the Glanville fritillary butterfly subject to highly fragmented landscapes (Hanski 2011, Hanski et al. 2017). In this unique example, environmental stochasticity makes dispersal profitable by creating new patches that can be colonized by dispersers. The prime importance of spatiotemporal patch variability in promoting dispersal is well supported by a number of theoretical models (Comins et al. 1980, McPeek & Holt 1992, Armsworth & Rougharden 2005), and is also advanced as the main driver of wing dimorphism observed in insects in a gradient of patch temporality (Denno et al. 1996).

Habitat alteration in human-dominated landscapes is often associated with shifts in disturbance regimes (Turner 2010, Newman 2019). For instance, one decade-long worldwide survey revealed a relatively weak net surface loss of temperate forests, but a high turnover due to forestry practices (Hansen et al. 2013). Shifts in the disturbance regimes prevailing in habitat remnants could therefore mitigate or, conversely, magnify the negative effect of habitat fragmentation, depending on their direction and magnitude. Despite this, apart for aerial dispersal in invertebrates (Denno et al. 1996), human-induced temporal variation in the spatial distribution of habitat patches has generally been overlooked when considering dispersal evolution in anthropogenic landscapes. A full appraisal of the effect of anthropogenic disturbance on dispersal should not only examine whether a dispersal pattern emerges in a landscape, but whether it gives rise to a genetic footprint throughout successive generations – and last but not least, to what extent this involves a specialized phenotype. Considering all the facets of this issue is not a simple task (Kokko & Lopez-Sepulcre 2006, Broquet & Petit 2009, Ronce & Clobert 2012), and to our knowledge has not yet been investigated in vertebrates.

To address this gap, this study examined how human-driven variation in habitat patch turnover affects dispersal rates and distances, determines dispersal-related phenotypic specialization, and drives neutral genetic variation in spatially structured populations. We studied this issue in an early successional amphibian, the yellow-bellied toad (*Bombina variegata*), a species that reproduces in small waterbodies with a short hydroperiod occurring either in (virtually) undisturbed or anthropogenic environments (Warren & Büttner 2008; Cayuela et al. 2011, 2015b). In riverine environments with negligible human activity, the species’ habitat patches are groups of rocky pools that result from long-term geomorphological processes alongside riverbanks (riverine SSPs; Cayuela et al. 2011). This results in a negligible patch turnover rate and makes patches available and predictable far beyond a toad’s lifespan. In contrast, in harvested woodlands, habitat patches consist of groups of ruts made by logging vehicles that may appear and disappear yearly as a result of the combined effects of logging operations and rapid natural silting (we refer to these as logging SSPs hereafter). This leads to a high patch turnover rate and makes patch location and availability more unpredictable at the scale of a toad’s lifespan (Cayuela et al. 2016a, 2016b). Previous studies have highlighted demographic differences in SSPs from the two environments and found that individuals in logging SSPs have a faster life history (i.e. a shorter lifespan and higher fecundity; Cayuela et al. 2016a), experience earlier senescence (Cayuela et al. 2019b), and display higher breeding dispersal probability (Cayuela et al. 2016b) than individuals in riverine habitats. In this study, our first step was to quantify dispersal probability and distance throughout an individual’s lifetime, as natal dispersal was lacking in previous studies and a review of recent literature suggested that natal and breeding dispersal patterns can strongly differ in amphibians (Cayuela et al. 2018c). We expected that (1) both natal and breeding dispersal rates and distances would be higher in logging than in riverine SSPs. In a second step, we analyzed how patch turnover and related dispersal costs and benefits affected SSP dynamics and long-term viability using simulations based on published demographic rates. We hypothesized that (2) dispersal and context-dependent immigration (i.e. depending on patch age) allows the long-term persistence of logging SSPs. In a third step, we used common garden experiments to investigate how patch turnover determines dispersal syndromes and may act as a selective agent on phenotypic specialization in riverine and logging SSPs. We expected (3) toadlets from logging SSPs to have behavioral traits (i.e. high exploration propensity and boldness) and morphological traits (i.e. long hind limbs) that generally facilitate dispersal in amphibians (reviewed in Cayuela et al. 2018c). In a fourth step, we examined how human-driven variation in patch turnover, by affecting neutral genetic diversity and gene flow, leads to contrasting genetic footprints over the longer term in riverine and logging SSPs. As genetic differentiation results from the combined effects of genetic drift and gene flow (Broquet & Petit 2009, Cayuela et al. 2018a), we selected two SSPs per landscape type in order to determine the relative contribution of each of these drivers. As high dispersal is expected to increase gene flow and decrease the local effects of genetic drift, we expected (4) a larger effective population size as well as a lower level of inbreeding and intra-patch relatedness in logging than in riverine SSPs. We also expected that (5) after controlling for SSP size and landscape connectivity, higher gene flow would lead to lower genetic structure and weaker genetic Isolation-By-Distance (IBD) pattern in logging than in riverine SSPs.

## Materials and methods

### Study area and sampled populations

The study was conducted in 8 SSPs in eastern France, three in riverine environments (R1, R2, and R3) and five in logging environments (L1, L2, L3, L4 and L5). The SSPs were chosen according to technical constraints or to minimize bias at each stage of the study – our choices are explained below. The distance separating SSPs from each other varied from 20–500 km (**Fig. S1**).

### Dispersal patterns throughout toad lifespan in riverine and logging environments

#### Studied populations

We quantified natal and breeding dispersal rates and distances in four SSPs (L1, L2 and R1, R2; see maps **Fig. S1** and **Fig. S2**) for which breeding rate dispersal had been previously estimated (Cayuela et al. 2016a, 2016b). The environmental characteristics of the four SSPs and the details regarding the survey design are presented in **Table 1** and **Table 2** respectively. The number of individuals captured each year is presented in **Supplementary material, Table S1**. A detailed description of the four SSPs can also be found in two previous studies (Cayuela et al. 2016a, 2016b). The number of patches (defined as a group of ruts) occupied by each SSP ranged from 8 to 189. Each SSP was monitored for a period of at least five years in one to five capture sessions per year that were usually between two weeks to one month apart. At each capture session, all the patches were sampled in the daytime and toads were captured by hand or dipnet. Based on previous studies (Cayuela et al. 2016a, 2016c), we considered three life stages: juveniles (i.e. post-wintering metamorphs), subadults (two-year-old immature animals) and adults (i.e. breeders, three years old or more). This resulted in a total dataset of 12,721 individual capture–recapture (CR) histories.

**Table 1.** Environmental characteristics of the four SSPs (L1, L2, R1 and R2): patch persistence over time, environment type (logging vs riverine), patch isolation (mean distance in meters between two pond networks, and associated variation coefficient) and patch size (mean number of ponds within a patch and associated variation coefficient).

| SSP | Patch persistence | Environment | Patch isolation | Patch size |
| --- | --- | --- | --- | --- |
| L1 | 1 to 10 years | logging activity | 2864.50 (57%) | 3.70 (70%) |
| L2 | 1 to 10 years | logging activity | 4424.64 (53%) | 5.64 (88%) |
| R1 | >30 years | natural erosion | 676.57 (61%) | 4.52 (105%) |
| R2 | >30 years | natural erosion | 501.62 (61%) | 4.71 (93%) |

**Table 2.** Survey design characteristics for the four SSPs (L1, L2, R1 and R2) considered in the study: study period, survey duration, number of capture sessions performed over the survey period, total number of captures, total number of individuals identified during the survey and total number of sampled patches (pond networks).

| SSP | Study period | Survey duration | Capture sessions | Number of captures | Number of individuals | Number of sampled patches |
| --- | --- | --- | --- | --- | --- | --- |
| L1 | 2000–2008 | 9 years | 29 | 953 | 445 | 28 |
| L2 | 2012–2016 | 5 years | 15 | 16477 | 12192 | 189 |
| R1 | 2010–2014 | 5 years | 13 | 4747 | 1003 | 14 |
| R2 | 2010–2014 | 5 years | 13 | 3984 | 769 | 8 |

#### Naïve dispersal kernels

We first estimated a dispersal kernel based on distances recorded during sampling using a lognormal distribution for each population and life stage (juvenile, subadult, and adult). This allowed us to visualize the form of the kernel from raw data before building complex CR models.

#### The structure of the multievent model

For the needs of our study, we extended the CR multi-event model proposed by Lagrange et al. (2014), which allows estimating survival (ϕ) and dispersal (ψ) in numerous sites. By omitting site identity and distinguishing between individuals that stay and individuals that move, this model circumvents the computational issues usually encountered in standard multi-site CR models when the number of sites is large (Lebreton et al. 2009). Lagrange’s model includes states that incorporate information about whether an individual is occupying at *t* the same site as the one occupied at *t* − 1 (‘S’= stayed) or not (‘M’= moved). The model also includes information about whether the individual was captured (‘+’) or not (‘o’) at *t* − 1 and *t*. Recently, Tournier et al. (2017) extended Lagrange’s model by breaking down dispersal (ψ) into distinct parameters of departure (ε) and arrival (α). This new parameterization allows the estimation of the proportion of individuals arriving in sites of different quality or located at different distances from the source site.

We adapted this parameterization for our study to consider states incorporating information about individual capture (‘+’ and ‘o’) at *t* − 1 and *t* as well as movement status. We also included states with information about the individual’s age class: juvenile (‘j’), subadult (‘s’) and adult (‘a’). Additionally, we incorporated information about the Euclidian distance covered by dispersers between the departure and arrival patch using three distance classes: ‘1’ = 100–800 m, ‘2’ = 800–1500 m, ‘3’ > 1500 m. This led to the consideration of 37 states in the model (**Fig.1** and **Fig.2**). For example, an individual +jS+ was captured at *t* − 1 and *t*, was a juvenile, and remained in the same patch between *t* − 1 and *t*. An individual +sM1+ was captured at *t* − 1 and *t*, was a subadult, did not occupy the same patch as at *t* − 1, and arrived in a patch located at a distance 100–800 m from the source patch. We distinguished 16 events, which were coded in an individual’s capture history and reflect the information available to the observer at the time of capture (**Fig.2**).

**Fig.1.**
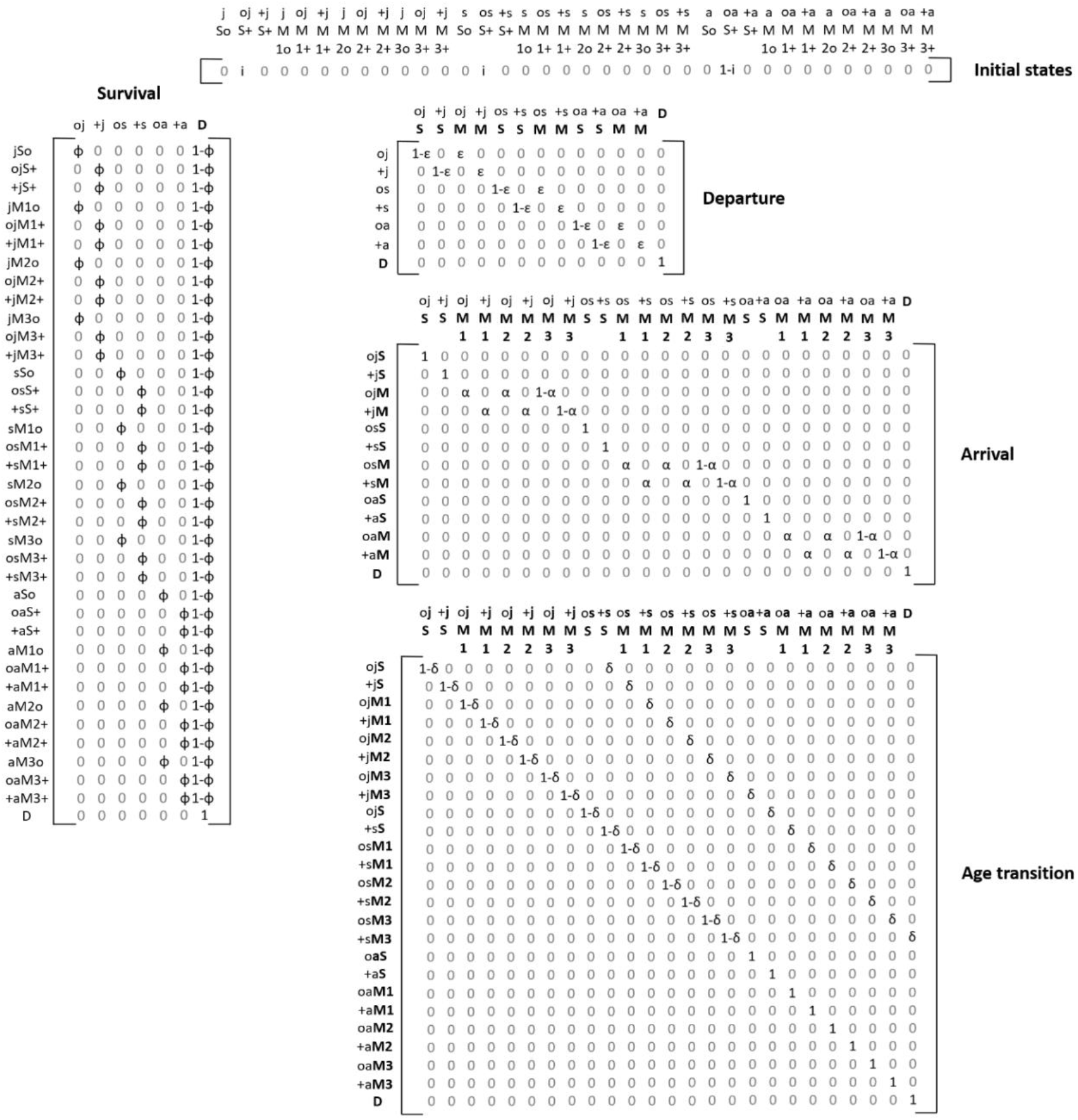
Model structure: matrices of initial states and state transitions (survival, departure, arrival and age transition). In the transition matrix, the rows correspond to time t − 1, the columns to time t, and whenever a status element is updated to its situation at t, it becomes bold and stays bold throughout the following steps.

When captured for the first time, the state of an individual could be ojS+, osS+ or oaS+. We then considered five modeling steps in which the information of the state descriptor was progressively updated: survival (ϕ), departure (ε), arrival (α), age transition (δ) and recapture (*p*). Each step was conditional on all previous steps. In the first step, we updated information about survival. An individual could survive with a probability of ϕ or die (D) with a probability of 1 − ϕ. This led to a matrix with 37 states of departure and 7 intermediate states of arrival (Fig.1). Survival probability could differ between age classes by allowing differing values for ϕ in lines 1–12, 13–24 and 25–36 of the matrix. In the second modeling step, departure was updated. Individuals could move (M) from the site they occupied with a probability of ε or stay (S) with a probability of 1 − ε. A matrix of 7 departure states and 13 arrival states was considered (Fig.2). Departure probability could differ between age classes by allowing differing values for ε in lines 1–2, 3–4 and 5–6 of the matrix. In the third step, we updated the arrival information. An individual that moved could arrive in a patch located in the first two distance classes (1 or 2) from the source patch with a probability of α, or arrive in a site located in the third distance class (3) with a probability of 1 − α. This resulted in a matrix with 13 departure states and 25 arrival states (Fig.2). Arrival probability could differ between age classes by allowing different values for α in lines 2–4, 7–8 and 11–12 of the matrix. In the fourth step, the information about age was updated. An individual could reach the next age class (j, s or a) with a probability of δ or remain in the previous age class with a probability of 1 − δ, resulting in a transition matrix with 25 states of departure and 25 states of arrival (**Fig.2**). The adult individuals (a) were forced to stay in their age class. In the fifth and last step, recapture was updated (**Fig.2**). An individual could be recaptured with a probability of *p* or missed with a probability of 1 − *p*, resulting in a transition matrix with 25 states of departure and 37 states of arrival. The recapture probability could differ between age classes by allowing different values for *p* in lines 1–18, 9–16 and 17–24 of the matrix. The last component of the model linked events to states. In this specific situation, each state corresponded to only one possible event (**Fig.2**).

**Fig.2.**
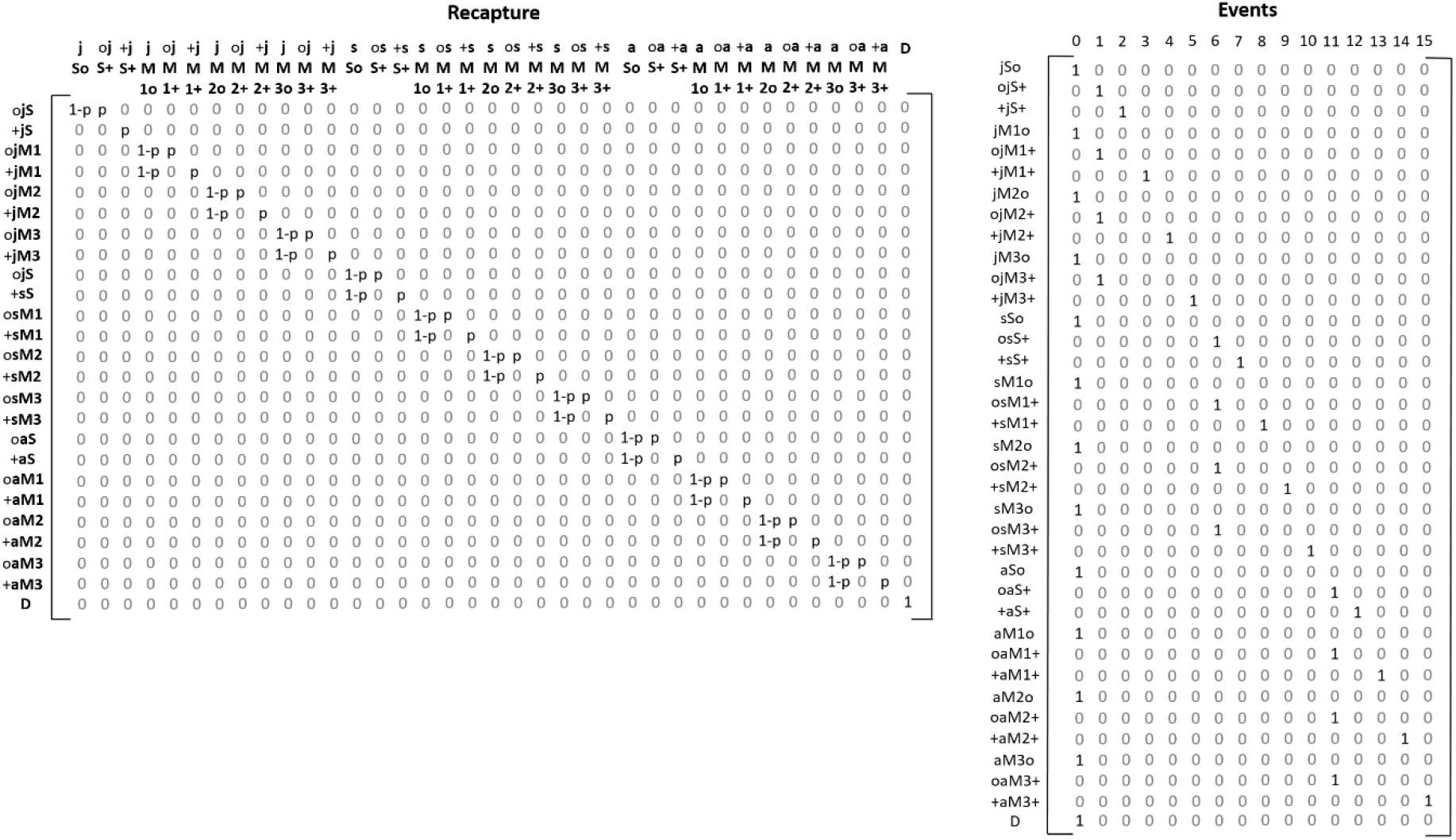
Model structure: state transitions (recapture) and events (field observations). In the transition matrix, the rows correspond to time t − 1, the columns to time t, and whenever a status element is updated to its situation at t, it becomes bold and stays bold throughout the following steps.

#### Biological scenarios in the E-SURGE program

The parameterization was implemented in the E-SURGE program (Choquet et al. 2009). The datasets for the four SSPs considered in our study were analyzed separately, as the number of study years and capture sessions for these populations varied. Competing models were ranked through a model-selection procedure using Akaike information criteria adjusted for a small sample size (AICc) and AICc weights. Following the recommendation of Burnham & Anderson, we performed model averaging when the AICc weight of the best-supported model was less than 0.90. The models had a robust design structure (Pollock 1982). As in previous studies of *B. variegata*, the survival probability was fixed at one between secondary sessions (Cayuela et al. 2016a, 2016c). The robust design structure allowed both intra-annual and inter-annual dispersal to be considered. Our hypotheses concerning recapture and state–state transition probabilities were tested using the general model [ϕ(AGE), ε(AGE), α(AGE), δ(.), *p*(AGE + Y)], which included two effects: (1) the three age classes (AGE) coded as states in the model, and (2) year-specific variation (Y). The notation (.) indicated that the parameter was held constant. We tested whether survival (ϕ) and departure probabilities (ε) varied between age class (AGE). Moreover, we hypothesized that the probability of arriving in a patch depends on age (AGE), and on the Euclidean distance between patches (the distance classes were incorporated as states in the model). Recapture probability was expected to differ between age classes (AGE) and years (Y). We tested our expectations about the model parameters in a stepwise fashion. From this general model, we tested all the possible combinations of effects and ran 16 competing models.

### Simulating the effect of patch turnover and dispersal on SSP dynamics and long-term persistence in logging contexts

We simulated population trajectories based on different scenarios to investigate the effects of patch turnover and dispersal costs on the dynamics and long-term persistence of SSPs in habitats subjected to logging. Adopting the most realistic lifecycle for the yellow-bellied toad (see ‘Results’, **Fig.4**) determined in previous studies (e.g. Cayuela et al. 2015, 2018), we used a three age-class (juveniles, subadults, and adults), female-dominant, prebreeding Leslie matrix (Caswell 2001) (see ‘Results’, **Fig.4**). We used the demographic parameters of a riverine SSP (R1; see Cayuela et al. 2016), which was considered a reference population whose demographic parameters have not been altered by dispersal costs (survival in logging SSPs is lower than in riverine SSPs, likely due to dispersal costs, Cayuela et al. 2018b). Both prebreeding survival probability (juvenile survival, S1 = 0.70; subadult survival, S2 = 0.77) and adult survival probability (S3 =0.92) were included in the Leslie matrix. Fecundity *F* was possible only for adult females and consisted of estimated recruitment: that is, the number of recruited juvenile females at *t* per breeding female at *t* − 1 (F = 0.52, Cayuela et al. 2016). As Boualit et al. (2019) found that juvenile recruitment was higher in newly created and disturbed patches than in old, undisturbed patches in logging SSPs, we specified that *F* decrease linearly (−5% per year) with the age of the patch. Furthermore, we also considered the possibility that females may skip breeding opportunities; in the riverine SSP R1, a previous study showed that the probability of females skipping breeding (B) was 0.15 (Cayuela et al. 2016). As in Cayuela et al. (2018, 2019c), we considered demographic stochasticity for survival, fecundity and skipping breeding. For each year, demographic parameter values were randomly sampled in a Gaussian distribution centered on mean parameter estimates, and standard deviation was inferred from two previous studies conducted on this species (Cayuela et al. 2016a, 2018b). The standard deviation values were: 0.05 for S1, 0.03 for S2, 0.01 for S3, 0.02 for *F*, and 0.02 for B.

Three patch turnover scenarios were considered. In scenario 1 (high turnover), a patch disappeared three years after its creation, in scenario 2 (medium turnover) a patch disappeared after six years, and in scenario 3 (low turnover) it disappeared after nine years. These scenarios correspond to the range of patch turnover in logging SSPs reported by forest managers (Eric Bonnaire, *unpublished data*), variation that depends on local management policies and the frequency of forest harvesting operations. We also considered three dispersal scenarios. In scenario 1 (no dispersal), individuals were not able to escape and died when a patch disappeared. In scenario 2 (dispersal with random immigration), individuals could disperse to escape the disappearance of a patch or could disperse by choice (i.e. when the patch remained available). The immigration was random between the patches of the metapopulation and was not influenced by the age of the patch. In scenario 3 (dispersal with informed immigration), individuals could disperse in response to patch disapperance or by choice. Based on an assessment of Boualit et al. (2019), we considered that immigration was not random and that immigration probability linearly decreases with patch age (i.e. a loss of 5% per year). In scenario 2 and 3, in which dispersal was possible, we considered two subsets of scenarios: in subset 1 (non-costly dispersal), individuals did not incur any survival loss when they dispersed. In subset 2 (costly dispersal), we considered that survival loss related to dispersal could be low (−5% of survival), medium (−10%), or high (−15%). As in Cayuela et al. (2018), we made the assumption that survival loss was similar across life stages (i.e. juvenile, subadult, and adult).

Each simulation began with 30 breeding patches over which 1,000 individuals were randomly scattered. The number of individuals in each age class was obtained through the stable stage distribution provided by the three age-class Leslie matrix. Then for each time step (a 1-year interval), we simulated the change in patch availability. We considered that five new patches were created each year. As the patches disappeared in a deterministic way when they reached the age defined in the scenario, the number of available patches remained constant over time (except for the few first years). We simulated the number of individuals in each age class occupying each patch. To do this, we separately considered patches reaching the age of disappearance versus those that did not disappear. In the latter, the number of individuals at *t* + 1 given the number of individuals at *t* was predicted by the Leslie matrix using the survival probability of individuals occupying an available patch (reported in the reference population R1). To be as realistic as possible, we used demographic stochasticity (fecundity was thus randomly sampled from a Poisson distribution, and survival from a binomial distribution), as in Cayuela et al. (2018). For patches that disappeared, we applied the same procedure, but using the survival probability (affected or not by dispersal cost, depending on the scenario) for individuals occupying patches that subsequently disappeared. Surviving individuals from lost patches were then randomly spread over the available patches at *t* + 1 in the ‘dispersal with random immigration’ scenario, or they were preferentially distributed in new patches in the ‘dispersal with informed immigration’ scenario. In all dispersal scenarios, we fixed the elective dispersal probability (dispersal when the patch did not disappear) *D* at 0.15, which was consistent with the annual dispersal rate reported in logging SSPs in the study. We also considered demographic stochasticity in elective dispersal (the standard deviation value was 0.05). The modeled population was monitored for 100 years. We did not remove the first few years of the simulation, when the number of patches progressively increased since none were old enough to disappear yet, as these had virtually no impact on our results (Cayuela et al. 2018). We performed 1,000 simulations for each scenario. At each time step, we monitored the number of adults in the entire SSP as well as the proportion of simulations in which the SSP went extinct (the extinction probability).

### Phenotypic specialization in riverine and logging environments

#### Study populations

To compare the morphology and behavior of toadlets in riverine SSPs with those of logging SSPs, we used a common garden experiment. This involved collecting between 8 and 15 egg clutches (hereafter referred to as ‘family’) in three SSPs in each landscape type (riverine = R1, R2, R3, and logging = L3, L4, L5; **Fig.S1**). These were selected to minimize spatial proximity between SSPs belonging to the same landscape type and therefore to avoid potential confounding effects resulting from spatial autocorrelation in environmental conditions. Five siblings per family were randomly chosen just after hatching and individually reared under controlled laboratory conditions.

#### Rearing protocol

The egg clutches were carefully transported to the laboratory, where they were individually placed in aquariums (32×17 cm, height 15 cm) with aged tap water equipped with an oxygen pump. The aquariums were placed in a climatic room with a light/dark cycle of 18:6, corresponding to the natural light/dark cycle in the area in the summer, with an ambient temperature varying from 21.5 to 23.5°C. Embryo development ended 2–5 days after the arrival of the egg clutches at the laboratory. After hatchling, when tadpoles reached Gosner stage 24–25 (active swimming, external gill atrophy) (Gosner 1960), five siblings per family were randomly chosen and placed individually into plastic containers (14×8.5 cm, height 13.5 cm) filled with previously aged tap water and aerated in a tank. The plastic containers were distributed in a predetermined random pattern around the climatically controlled room. The water was replaced every three days. The tadpoles were fed every day with 150 mg of cooked lettuce, providing ad libitum feeding. At Gosner stage 44–45 (tail atrophy, mouth posterior to eyes), feeding was terminated, and the water was drained and replaced with a dampened sponge placed in the bottom of the container. The sponge and the walls were sprayed with aged tap water every two days. The individuals were kept until their complete metamorphosis (Gosner stage 46) and were then subjected to behavioral assays.

#### Experimental arenas

The behavioral assays took place in an arena (70 cm in diameter) made of polyethylene terephthalate, with a central shelter and a ‘desiccation’ obstacle between this and the 12 possible exits (**Fig. S4**). At the center of the arena, we placed a removable, cylindrical (9 cm diameter, height 90 cm), opaque, covered chamber (‘refuge chamber’ hereafter). The cylinder had a circular opening (3 cm diameter) covered by a lid. Around the interior edge of the arena, we installed a pit (width 10 cm, depth 0.3 cm) filled with a desiccating mixture of sand and highly active silica gel powder in a weight ratio of 0.8:0.2 (‘desiccant zone’ hereafter). The arena wall included 12 doors placed at regular intervals around the entire circumference. The arena was confined in an enclosed iron chamber (150×17 cm, height 175 cm) over which a dark opaque sheet was placed to limit potential acoustic and visual interference during behavioral trials.

#### Behavioral assays

At metamorphosis, each toadlet was subjected to a behavioral assay to quantify their neophobia or exploratory behavior. The behavioral tests were conducted in the circular arena described above. Before each trial, the toadlet and a dampened sponge (already present in the toadlet’s rearing container) were gently transferred into an opaque circular release box that was then placed at the center of the arena. The dampened sponge was considered a known object, making the refuge chamber more familiar than the rest of the experimental device. Following previous studies on anuran behavioral syndromes (reviewed in Kelleher et al. 2018), neophobia was quantified as the latency time to enter a novel environment (i.e. the time delay to leave the familiar refuge chamber: BEHAV1). Exploration propensity was assessed using two variables: the latency time to enter a novel but harsh environment (i.e. to reach the desiccant zone after leaving the refuge chamber; BEHAV2), and the latency time to travel the harsh environment and get out of the arena (BEHAV3). The behavior was recorded for 30 min (1800 s) using a digital camera (Sony DCR-SX34).

#### Extraction of behavioral variables from the videos

The videos were analysed using the BEMOVI R package (Pennekamp et al. 2015) to reconstruct the movement trajectory of the toadlet in the arena, and to extract a series of behavioral variables from this. First, the videos were standardized to a length of 29 min (the minimum duration available for all individuals) removing the initial 60 s and the extra time at the end, if any. Then the videos were converted to a format suitable for analysis in BEMOVI: rectangular pixels (720 x 576) were converted into square pixels (1024 x 576), color information was converted into 256 gray levels, frames per s were decreased from 25 to 5 to limit memory allocation requirements, and the videos were saved as AVI files. These operations were performed using FFMPEG software (ffmpeg.org).

BEMOVI was then run with the following parameters: black and white threshold (40) to discriminate the toadlet from the background; minimum size (20) and maximum size (150), corresponding to the size range of the toadlets; link range (7500 frames) to allow any duration of the ‘disappearance’ of the toadlet from the video (e.g. when it was in the refuge) while still considering it as a single movement trajectory, and “disp “(100 pixels); in BEMOVI, disp is the maximal distance covered by the toadlet from one frame to the next, corresponding to 0.2s here. This resulted in a database with the movement trajectory of each toadlet, i.e. its X–Y position at each time step. This position was compared to the distance from the center of the arena to determine which zone (refuge chamber, normal zone or desiccant zone) the toadlet was in at each time step. The results were checked for errors in toadlet positioning, which were due to varying light conditions, usually at the beginning of the videos. In a final step, we computed several behavioral variables for each toadlet from this positioning information.

#### Statistical analyses

We used linear mixed models to test whether morphological traits (body size, body condition and relative hind limb size) of toadlets differed according to the landscape type of origin (logging vs riverine). Each morphological trait was treated as a dependent variable, and the landscape type was introduced as a fixed explanatory term in the model. In the case of body condition and relative leg size, body size and its interactive effect with the landscape type were also introduced as adjustment covariates in the fixed part of the model. Both the SSP of origin and the clutch were introduced as random effects in the model. We also allowed heterogeneity of variance between landscape types by allowing a separate estimation of the residual variance for each landscape type. The estimation method was based on restricted maximum likelihood. Variance heterogeneity was first checked using the likelihood ratio test and removed if non-significant. The significance of each explanatory term was examined with a non-sequential *F* test based on the Kenward-Roger method to approximate the denominator *df* (Littel et al. 2006). In the case of covariance analyses (body condition and relative leg size), the interactive effect was discarded if non-significant to obtain the final model. All morphological variables were standardized using Z transformation before the analyses, as recommended by Schielzeth (2010).

We tested whether the behavioral variables BEHAV1 and BEHAV2 varied according to the landscape type (i.e. logging vs riverine) using a generalized linear mixed model. Each latency time variable was treated as a dependent variable using a Poisson distribution. The landscape type was introduced as an explanatory term in the fixed part of the model. We also introduced individual body size and its interactive effect with the landscape of origin as adjustment covariates in the fixed part of the model since both locomotion skill and exploratory performance can vary according to individual size (reviewed in Kelleher et al. 2018). For the analyses of morphological traits, both the SSP of origin and the clutch were introduced in the model as random effects. Furthermore, a scale parameter was also introduced to handle data overdispersion and to obtain a corrected statistical test using a quasi-likelihood approach (McCullagh & Nelder 1989). The estimation method was based on restricted pseudo-likelihood optimization, and the significance of each explanatory term was examined using the same methodology as for the morphological analyses. Non-significant terms were successively removed to obtain the final model, and least square means were used to estimate the difference in latency time variables according to the landscape of origin.

As the third behavioral variable (BEHAV3) was right censored, it was analyzed using a proportional hazards mixed-effects model (i.e. a frailty model based on a Cox model, PHREG procedure, SAS Institute Inc. 2012). We tested whether newborn individuals originating from logging systems were more prone to exit the assay arena than those from riverine systems. It is not possible to handle multiple random factors in such a model, so we took into account only the family effect since this was found to be significant for the other behavioral variables but not for the SSP of origin. The landscape of origin, the body size and their interactive effect were introduced as explanatory terms. Parameters were estimated using partial likelihood estimation, and the significance of explanatory terms was assessed using non-sequential chi-square tests.

### Neutral genetic variation in riverine and logging environments

#### Study populations

We examined neutral genetic variations within two SSPs in riverine environments (R1 and R2) and two SSPs in logging environments (L3 and L4) (see map in **Fig. S3**) using 15 polymorphic microsatellite markers (described and tested in Cayuela et al. 2017a). The four SSPs were selected according to the following criteria: (1) SSPs embedded in a relatively continuous forested matrix to avoid any confounding effect of matrix composition on gene flow (**Fig. S3**) – woodland is generally considered highly favorable for the movement of forest amphibians (Cushman 2006a) such as *B. variegata* (Cayuela et al. 2015b); and (2) two small (R1 and L3) and two large SSPs (R2 and L4) to control for genetic drift. The number of patches and DNA sampled per SSP are given in **Table S2**. We used the protocol described in Cayuela et al. (2017) for DNA extraction and amplification, individual genotyping, and allele scoring.

#### Estimating basic genetic metrics

We examined basic assumptions (i.e. detection of null alleles, Hardy-Weinberg equilibrium) and descriptive statistics (mean number of alleles per locus *N_a_*; effective number of alleles per locus *N_ef_*; observed heterozygosity *H_0_*; and expected heterozygosity *H_e_*) of the genetic diversity within each SSP. *N_a_*, *N_ef_*, *H_o_* and *H_e_* were calculated using GENALEX (Peakall & Smouse 2006). The inbreeding coefficient *F_is_* and the Hardy–Weinberg deviation were assessed with GENEPOP 4.1 (Rousset 2008). The detection of null alleles was performed using the program MICRO-CHECKER 2.2.3 (Van Oosterhout et al. 2004). Two microsatellite markers, Bomvar_Cons470 and bv11.7, were discarded from the subsequent analyses due to the presence of null alleles in the four SSPs. All descriptive statistics are provided in **Table S3**.

#### Estimating relatedness, inbreeding, and effective population size

We estimated relatedness and individual inbreeding using COANCESTRY v1.0.1.8 (Wang 2011). We performed simulations to identify the best relatedness and inbreeding estimator for our combined datasets; these consisted of 1,000 dyads spread equally across six categories of relatedness: parent–offspring (rxy = 0.5), full siblings (rxy = 0.5), half siblings/avuncular/grandparent–grandchild (rxy = 0.25), first cousins (rxy = 0.125), second cousins (rxy = 0.03125), and unrelated (rxy = 0). According to our simulations, the best rxy estimator for relatedness analysis was DyadML, which showed a strong correlation of 0.74. We used linear mixed models to evaluate how SSP size (small: R1 and L4; large: R2 and L3) and the type of environment (logging vs riverine) affected individual inbreeding and intra-patch relatedness. An SSP’s size and environment type were coded as fixed effects (we considered an interaction between the two factors), whereas patch identity was coded as a random effect. We then estimated the effective population size of the four SSPs using the linkage disequilibrium method implemented in Ne Estimator v2.1 (Do et al. 2014).

#### Clustering approach

We described the genetic differentiation between patches within each SSP using an assignation method based on the Bayesian clustering algorithm implemented in the software STRUCTURE (Pritchard et al. 2000). Specifically, we estimated the most likely number of genetic clusters (K) contained in each SSP following the hierarchical approach proposed by Balkenhol et al. (2014) to detect additional substructures within clusters. The STRUCTURE program was run with the admixture model, with a burn-in period of 100,000 repetitions, and 100,000 subsequent MCMC repetitions. The *K* values were tested ranging from 1 to 10 and analyses repeated 10 times for each value. We used STRUCTURE HARVESTER (Earl 2012) to summarize the results, determining the optimal *K* value using both log-likelihood plots and the delta *K* statistic (Evanno et al. 2005). We followed the hierarchical approach proposed by Coulon et al. (2008) to test for additional population substructures within clusters. Accordingly, these analyses were then repeated for each inferred population cluster separately until the optimal *K* value was 1 (meaning that no additional structure was found within clusters). To map the spatial distribution of the different clusters, the individual ancestry values were averaged across the ten STRUCTURE runs using CLUMMP software (Jakobsson & Rosenberg 2007). The ‘greedy’ algorithm in the CLUMMP program was then used to assign the individuals to the cluster in which they showed the highest Q-values.

#### Gradient analyses

We used direct gradient analyses (Prunier et al. 2015) to test whether spatial genetic differentiation was lower within logging than riverine SSPs while controlling for potential differences in the functional connectivity prevailing within each SSP. To do this, pairwise genetic distances between all individuals from each SSP were computed using the Bray-Curtis percentage dissimilarity measures (Cushman et al. 2006b, Legendre & Legendre 2012), and were then standardized separately for each SSP. Since by design all SSPs were embedded in a relatively homogeneous forested matrix, we did not control for the effect of land cover type in these analyses. Rather, we focused on the effect of topographic roughness (‘slope’ effect) and the hydrological network topology (‘network’ effect) since both these landscape features have often been reported to affect gene flow in amphibians (Lowe 2003, Grant et al. 2010).

For each SSP, resistance layers were produced using ArcGIS Pro. Minimum geographical bounding of each SSP was determined and extended to a distance of five kilometres to avoid any edge effects. The elevation raster and hydrographic network shapefile, available through the French National Institute for Geographical and Forest Information database (BD ALTI®, and BD TOPO®), were extracted for SSP surface areas. To determine the percentage of steepness raster maps, the slope tool was used. Due to the spatial resolution imposed by elevation data, all layers were converted to raster format and homogenized at a spatial resolution of five meters.

Four resistance maps were thus constructed for each SSP. The first map included only the effect of geographic distance (i.e. isolation by distance, hereafter geographic resistance); it was based on data extended to include a 5-km buffer and was assigned a uniform resistance of 1 unit for all pixels. The second map included the topographic roughness (slope), where the resistance of each pixel was a linear function of the steepness (resistance = 1 + degree of steepness), thus corresponding to the effect of both geographic distance and topographic roughness. The third map included the hydrological network topology (network), where the resistance was assigned to 1 unit for all pixels situated in the hydrological network and to 10 units otherwise. The fourth map included both the slope effect and the network effect. Pairwise resistance between all patches was then computed from these four resistance maps using the circuit theory in Circuitscape V4.0 (McRae & Beier 2007).

Linear mixed models for pairwise distance matrices were then used to assess the effect of the geographic distance between patches (i.e. the geographic resistance) on the genetic distance between toads within each SSP while controlling for the effect of topographic roughness and/or the hydrological network topology. The non-independence between pairwise distances was taken into account in the covariance structure of the models. Specifically, we used the method proposed by Clarke et al. (2002) using a Toeplitz(1) as a covariance structure to specify the non-independence of pairwise genetic distance according to the patches of origin (see Selkoe et al. 2010, van Strien et al. 2012 for application on pairwise Fst distances). We used the extension presented in Prunier et al. (2013) for application on individual genetic distance in spatially hierarchized sampling schemes as is the case in our study. This resulted in two covariance parameters for each SSP, one for the patch random effect and the other for the individual random effect. Since both slope and network resistance were highly correlated to geographic resistance and as our main aim was to estimate the effect of geographic resistance on genetic distance while controlling for the effect of functional connectivity, we first regressed each of these effective resistances on each geographic resistance using simple linear regressions to obtain uncorrelated effective resistance respectively related to the slope effect, the network effect and their combined effect. The relative validity of each alternative landscape hypothesis (i.e. including the effect of the slope or of the stream network or both or neither on the genetic distances) was evaluated using weighted AIC (Waic), and the model-averaged estimate of the beta weight associated with the effect of geographic resistance on genetic distance was computed for each SSP.

## Results

### Dispersal patterns throughout toad lifespan in riverine and logging environments

Model averaging estimates (see **Table S4** for survival and recapture probabilities) indicated that both natal and breeding dispersal rates were high in logging environments (around 20% per year, **Fig.3**). In contrast, in riverine environments, natal dispersal rates were null and breeding dispersal rates were very low (less than 5% per year, **Fig.3**). This higher dispersal propensity in logging than in riverine habitats is even more remarkable given that the mean inter-patch distance is three times farther in the former than the latter (**Table S1**). The few adults that did disperse in riverine environments covered shorter distances: the median distance was 168 m in R1 and 189 m in R2, while the maximal distance was 455 m and 378 m respectively. This result was further confirmed by multi-event CR models showing that 100% of dispersal occurred over distances ranging from 100 to 800 m in riverine SSPs. In logging environments, dispersers covered substantially longer distances: the median distance was 431 m in L1 and 568 m in L2, while the maximal distance was 3810 m and 4529 m respectively. This result was supported by both the observed dispersal kernel and the multi-event model estimates for each life stage (see **Fig.3**). While these results indicate extremely contrasting dispersal regimes between logging and riverine landscapes, we also found substantial variations in the dispersal patterns in the two logging systems. First, while the inter-annual dispersal rate was similar in both logging systems, the intra-annual dispersal rate was substantially higher in L1 than L2 (i.e. L1 > 10 % and L2 <5%, **Fig.3**; for model selection procedure see **Table 3** and **Table 4**). Second, the dispersal kernels show a clear leptokurtic distribution decreasing with age in L2, suggesting a large demographic weight of natal dispersal in this SSP. This was not the case in L1, in which leptokurtosis was more reduced in juveniles than in adults.

**Fig.3.**
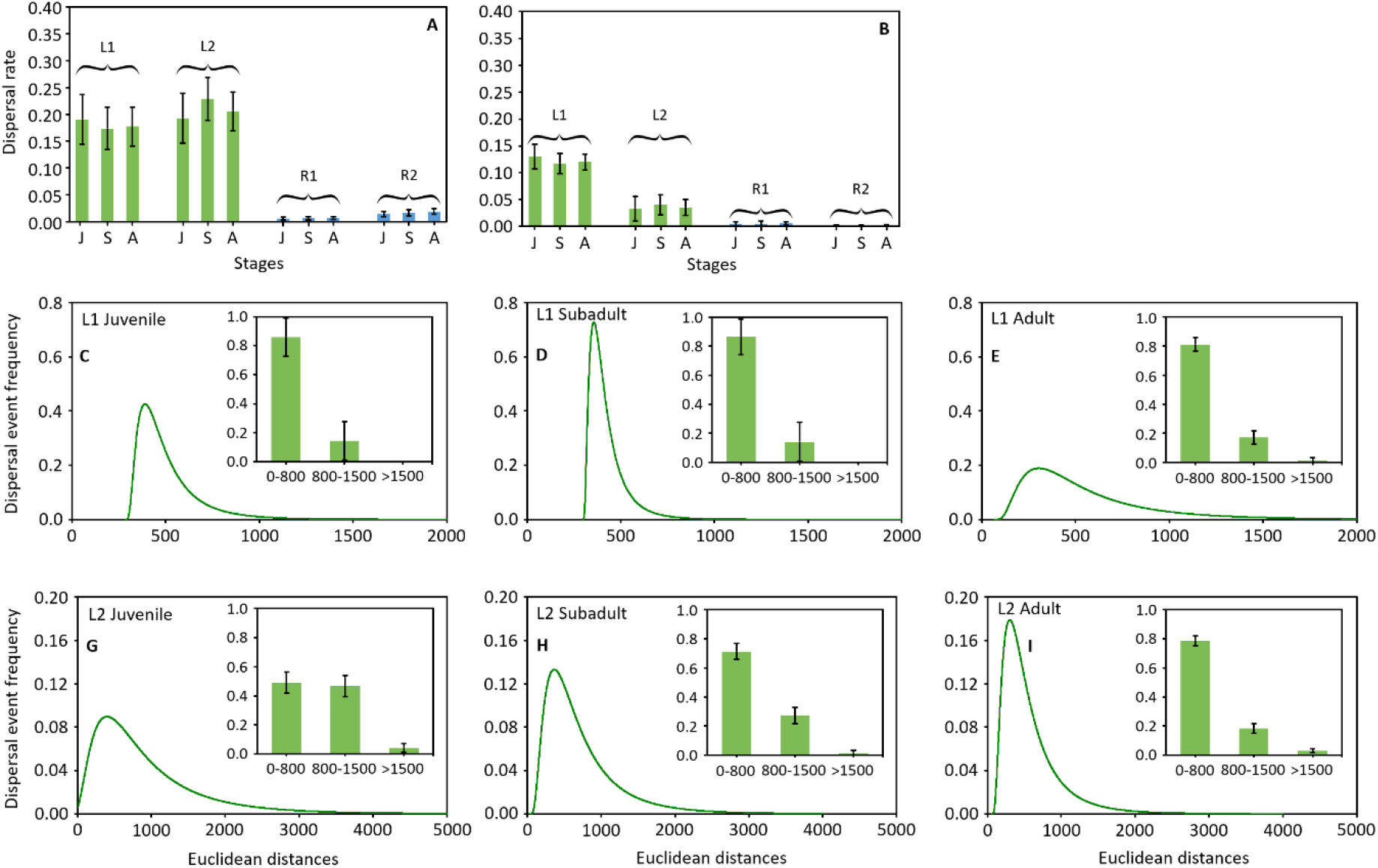
Natal and breeding dispersal rates and distances in logging (L1 and L2) and riverine SSPs (R1 and R2) (A–B). Inter-annual (A) and intra-annual (B) dispersal probabilities are higher in SSPs in logging landscapes than in riverine SSPs, regardless of the ontogenetic stage (juvenile, J; subadult, S; adult, A). Natal and breeding dispersal distances in logging SSPs (L1 and L2) (C–I). Dispersal event frequency decreases with the Euclidean distance between breeding patches at the ontogenetic stages.

**Table 3.**
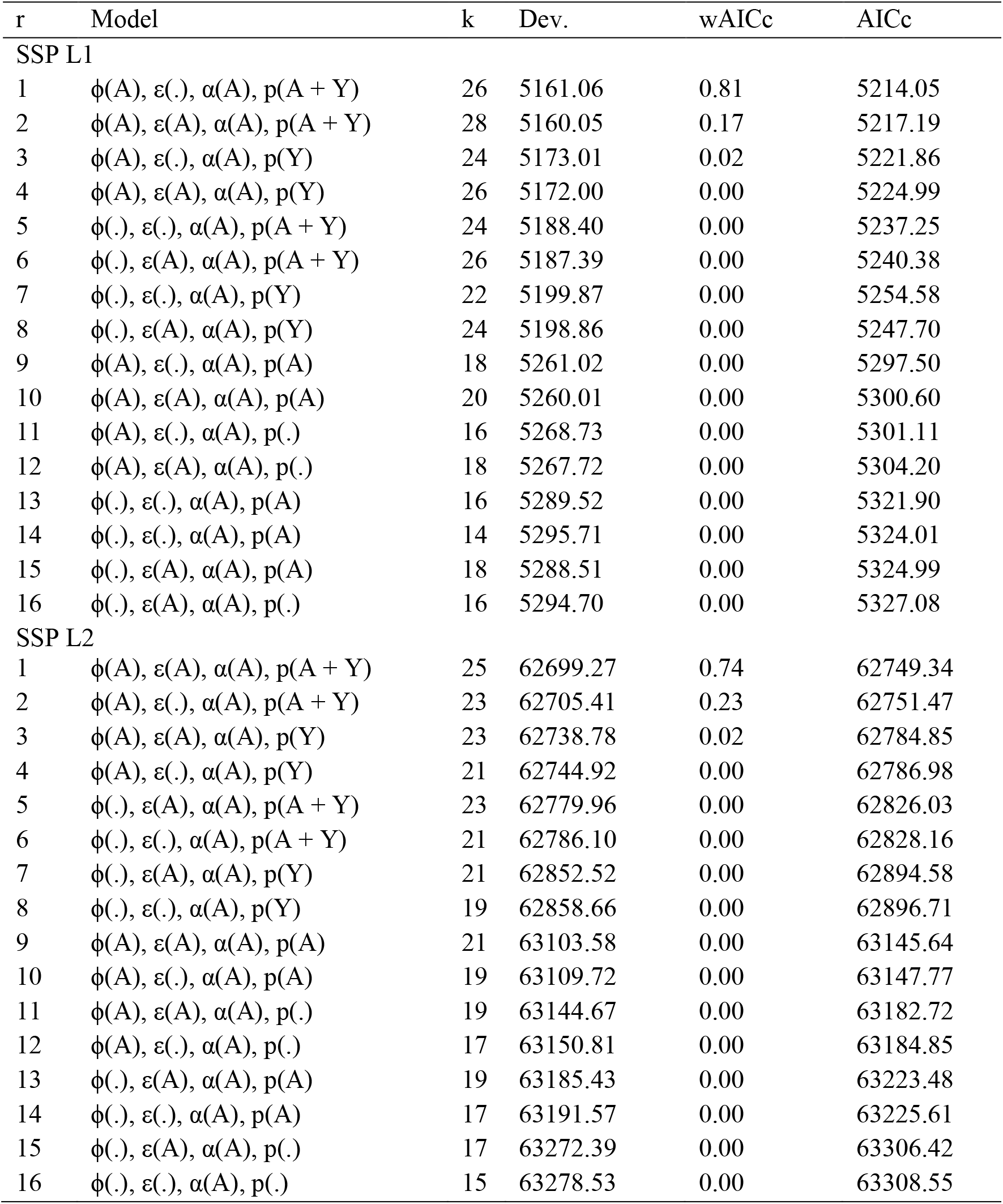
Multievent models and selection procedure for SSPs L1 and L2. r = model rank, k = number of parameters, Dev. = residual deviance, AICc = Akaike information criterion adjusted for small sample size, wAICc = AICc weight ϕ = survival probability, ε = departure probability, α = arrival probability, p = recapture probability, A = age,. = constant, Y = year.

| r | Model | k | Dev. | wAICc | AICc |
| --- | --- | --- | --- | --- | --- |
| SSP L1 |  |  |  |  |  |
| 1 | $\phi(A), \varepsilon(.), \alpha(A), p(A + Y)$ | 26 | 5161.06 | 0.81 | 5214.05 |
| 2 | $\phi(A), \varepsilon(A), \alpha(A), p(A + Y)$ | 28 | 5160.05 | 0.17 | 5217.19 |
| 3 | $\phi(A), \varepsilon(.), \alpha(A), p(Y)$ | 24 | 5173.01 | 0.02 | 5221.86 |
| 4 | $\phi(A), \varepsilon(A), \alpha(A), p(Y)$ | 26 | 5172.00 | 0.00 | 5224.99 |
| 5 | $\phi(.), \varepsilon(.), \alpha(A), p(A + Y)$ | 24 | 5188.40 | 0.00 | 5237.25 |
| 6 | $\phi(.), \varepsilon(A), \alpha(A), p(A + Y)$ | 26 | 5187.39 | 0.00 | 5240.38 |
| 7 | $\phi(.), \varepsilon(.), \alpha(A), p(Y)$ | 22 | 5199.87 | 0.00 | 5254.58 |
| 8 | $\phi(.), \varepsilon(A), \alpha(A), p(Y)$ | 24 | 5198.86 | 0.00 | 5247.70 |
| 9 | $\phi(A), \varepsilon(.), \alpha(A), p(A)$ | 18 | 5261.02 | 0.00 | 5297.50 |
| 10 | $\phi(A), \varepsilon(A), \alpha(A), p(A)$ | 20 | 5260.01 | 0.00 | 5300.60 |
| 11 | $\phi(A), \varepsilon(.), \alpha(A), p(.)$ | 16 | 5268.73 | 0.00 | 5301.11 |
| 12 | $\phi(A), \varepsilon(A), \alpha(A), p(.)$ | 18 | 5267.72 | 0.00 | 5304.20 |
| 13 | $\phi(.), \varepsilon(.), \alpha(A), p(A)$ | 16 | 5289.52 | 0.00 | 5321.90 |
| 14 | $\phi(.), \varepsilon(.), \alpha(A), p(A)$ | 14 | 5295.71 | 0.00 | 5324.01 |
| 15 | $\phi(.), \varepsilon(A), \alpha(A), p(A)$ | 18 | 5288.51 | 0.00 | 5324.99 |
| 16 | $\phi(.), \varepsilon(A), \alpha(A), p(.)$ | 16 | 5294.70 | 0.00 | 5327.08 |
| SSP L2 |  |  |  |  |  |
| 1 | $\phi(A), \varepsilon(A), \alpha(A), p(A + Y)$ | 25 | 62699.27 | 0.74 | 62749.34 |
| 2 | $\phi(A), \varepsilon(.), \alpha(A), p(A + Y)$ | 23 | 62705.41 | 0.23 | 62751.47 |
| 3 | $\phi(A), \varepsilon(A), \alpha(A), p(Y)$ | 23 | 62738.78 | 0.02 | 62784.85 |
| 4 | $\phi(A), \varepsilon(.), \alpha(A), p(Y)$ | 21 | 62744.92 | 0.00 | 62786.98 |
| 5 | $\phi(.), \varepsilon(A), \alpha(A), p(A + Y)$ | 23 | 62779.96 | 0.00 | 62826.03 |
| 6 | $\phi(.), \varepsilon(.), \alpha(A), p(A + Y)$ | 21 | 62786.10 | 0.00 | 62828.16 |
| 7 | $\phi(.), \varepsilon(A), \alpha(A), p(Y)$ | 21 | 62852.52 | 0.00 | 62894.58 |
| 8 | $\phi(.), \varepsilon(.), \alpha(A), p(Y)$ | 19 | 62858.66 | 0.00 | 62896.71 |
| 9 | $\phi(A), \varepsilon(A), \alpha(A), p(A)$ | 21 | 63103.58 | 0.00 | 63145.64 |
| 10 | $\phi(A), \varepsilon(.), \alpha(A), p(A)$ | 19 | 63109.72 | 0.00 | 63147.77 |
| 11 | $\phi(A), \varepsilon(A), \alpha(A), p(.)$ | 19 | 63144.67 | 0.00 | 63182.72 |
| 12 | $\phi(A), \varepsilon(.), \alpha(A), p(.)$ | 17 | 63150.81 | 0.00 | 63184.85 |
| 13 | $\phi(.), \varepsilon(A), \alpha(A), p(A)$ | 19 | 63185.43 | 0.00 | 63223.48 |
| 14 | $\phi(.), \varepsilon(.), \alpha(A), p(A)$ | 17 | 63191.57 | 0.00 | 63225.61 |
| 15 | $\phi(.), \varepsilon(A), \alpha(A), p(.)$ | 17 | 63272.39 | 0.00 | 63306.42 |
| 16 | $\phi(.), \varepsilon(.), \alpha(A), p(.)$ | 15 | 63278.53 | 0.00 | 63308.55 |

**Table 4.** Multievent models and selection procedure for SSPs R1 and R2. r = model rank, k = number of parameters, Dev. = residual deviance, AICc = Akaike information criterion adjusted for small sample size, wAICc = AICc weight ϕ = survival probability, ε = departure probability, α = arrival probability, p = recapture probability, A = age,. = constant, Y = year.

| r | Model | k | Dev. | wAICc | AICc |
| --- | --- | --- | --- | --- | --- |
| SSP R1 |  |  |  |  |  |
| 1 | $\phi(A), \varepsilon(A), \alpha(A), p(Y)$ | 19 | 8044.47 | 0.47 | 8082.67 |
| 2 | $\phi(A), \varepsilon(A), \alpha(A), p(A + Y)$ | 21 | 8041.98 | 0.22 | 8084.22 |
| 3 | $\phi(A), \varepsilon(\cdot), \alpha(A), p(Y)$ | 17 | 8050.14 | 0.21 | 8084.29 |
| 4 | $\phi(A), \varepsilon(\cdot), \alpha(A), p(A + Y)$ | 19 | 8047.65 | 0.10 | 8085.84 |
| 5 | $\phi(\cdot), \varepsilon(A), \alpha(A), p(Y)$ | 17 | 8074.54 | 0.00 | 8108.70 |
| 6 | $\phi(\cdot), \varepsilon(\cdot), \alpha(A), p(Y)$ | 15 | 8080.20 | 0.00 | 8110.32 |
| 7 | $\phi(\cdot), \varepsilon(A), \alpha(A), p(A + Y)$ | 19 | 8073.47 | 0.00 | 8111.67 |
| 8 | $\phi(\cdot), \varepsilon(\cdot), \alpha(A), p(A + Y)$ | 17 | 8079.13 | 0.00 | 8113.29 |
| 9 | $\phi(A), \varepsilon(A), \alpha(A), p(\cdot)$ | 15 | 8129.11 | 0.00 | 8159.23 |
| 10 | $\phi(A), \varepsilon(A), \alpha(A), p(A)$ | 17 | 8125.73 | 0.00 | 8159.89 |
| 11 | $\phi(A), \varepsilon(\cdot), \alpha(A), p(\cdot)$ | 13 | 8134.77 | 0.00 | 8160.87 |
| 12 | $\phi(A), \varepsilon(\cdot), \alpha(A), p(A)$ | 15 | 8131.39 | 0.00 | 8161.52 |
| 13 | $\phi(\cdot), \varepsilon(A), \alpha(A), p(\cdot)$ | 13 | 8157.05 | 0.00 | 8183.14 |
| 14 | $\phi(\cdot), \varepsilon(\cdot), \alpha(A), p(\cdot)$ | 11 | 8162.71 | 0.00 | 8184.78 |
| 15 | $\phi(\cdot), \varepsilon(A), \alpha(A), p(A)$ | 15 | 8155.63 | 0.00 | 8185.75 |
| 16 | $\phi(\cdot), \varepsilon(\cdot), \alpha(A), p(A)$ | 13 | 8161.29 | 0.00 | 8187.38 |
| SSP R2 |  |  |  |  |  |
| 1 | $\phi(A), \varepsilon(A), \alpha(A), p(A + Y)$ | 21 | 11041.21 | 0.64 | 11083.41 |
| 2 | $\phi(A), \varepsilon(\cdot), \alpha(A), p(A + Y)$ | 19 | 11047.74 | 0.19 | 11085.90 |
| 3 | $\phi(A), \varepsilon(A), \alpha(A), p(A)$ | 17 | 11052.43 | 0.13 | 11086.56 |
| 4 | $\phi(A), \varepsilon(\cdot), \alpha(A), p(A)$ | 15 | 11058.96 | 0.04 | 11089.06 |
| 5 | $\phi(\cdot), \varepsilon(A), \alpha(A), p(A + Y)$ | 19 | 11068.13 | 0.00 | 11106.29 |
| 6 | $\phi(\cdot), \varepsilon(\cdot), \alpha(A), p(A + Y)$ | 17 | 11074.66 | 0.00 | 11108.78 |
| 7 | $\phi(\cdot), \varepsilon(A), \alpha(A), p(A)$ | 15 | 11078.84 | 0.00 | 11108.94 |
| 8 | $\phi(\cdot), \varepsilon(\cdot), \alpha(A), p(A)$ | 13 | 11085.36 | 0.00 | 11111.44 |
| 9 | $\phi(A), \varepsilon(A), \alpha(A), p(Y)$ | 19 | 11076.17 | 0.00 | 11114.33 |
| 10 | $\phi(A), \varepsilon(\cdot), \alpha(A), p(Y)$ | 17 | 11082.69 | 0.00 | 11116.82 |
| 11 | $\phi(A), \varepsilon(A), \alpha(A), p(\cdot)$ | 15 | 11087.00 | 0.00 | 11117.10 |
| 12 | $\phi(A), \varepsilon(\cdot), \alpha(A), p(\cdot)$ | 13 | 11093.52 | 0.00 | 11119.60 |
| 13 | $\phi(\cdot), \varepsilon(A), \alpha(A), p(Y)$ | 17 | 11111.72 | 0.00 | 11145.85 |
| 14 | $\phi(\cdot), \varepsilon(\cdot), \alpha(A), p(Y)$ | 15 | 11118.24 | 0.00 | 11148.34 |
| 15 | $\phi(\cdot), \varepsilon(A), \alpha(A), p(\cdot)$ | 13 | 11122.54 | 0.00 | 11148.62 |
| 16 | $\phi(\cdot), \varepsilon(\cdot), \alpha(A), p(\cdot)$ | 11 | 11129.07 | 0.00 | 11151.12 |

### Simulating the effect of patch turnover and dispersal on logging SSP dynamics and long-term persistence

The results showed that patch turnover rate was a critical driver of logging SSP dynamics. The absence of dispersal within an SSP experiencing patch turnover necessarily led to the extinction of the SSP (**Fig.4A, 4B**, and **4C**), and the extinction speed increased with patch turnover rate. In contrast, patch turnover had a positive effect on SSP size (i.e. number of adults) when dispersal was possible and had no survival cost. This was caused by a fecundity-related mechanism: high patch turnover led to a decrease in the mean patch age within the SSP, resulting in an increase in average fecundity due to the positive relationship between fecundity and patch age.

**Fig.4.**
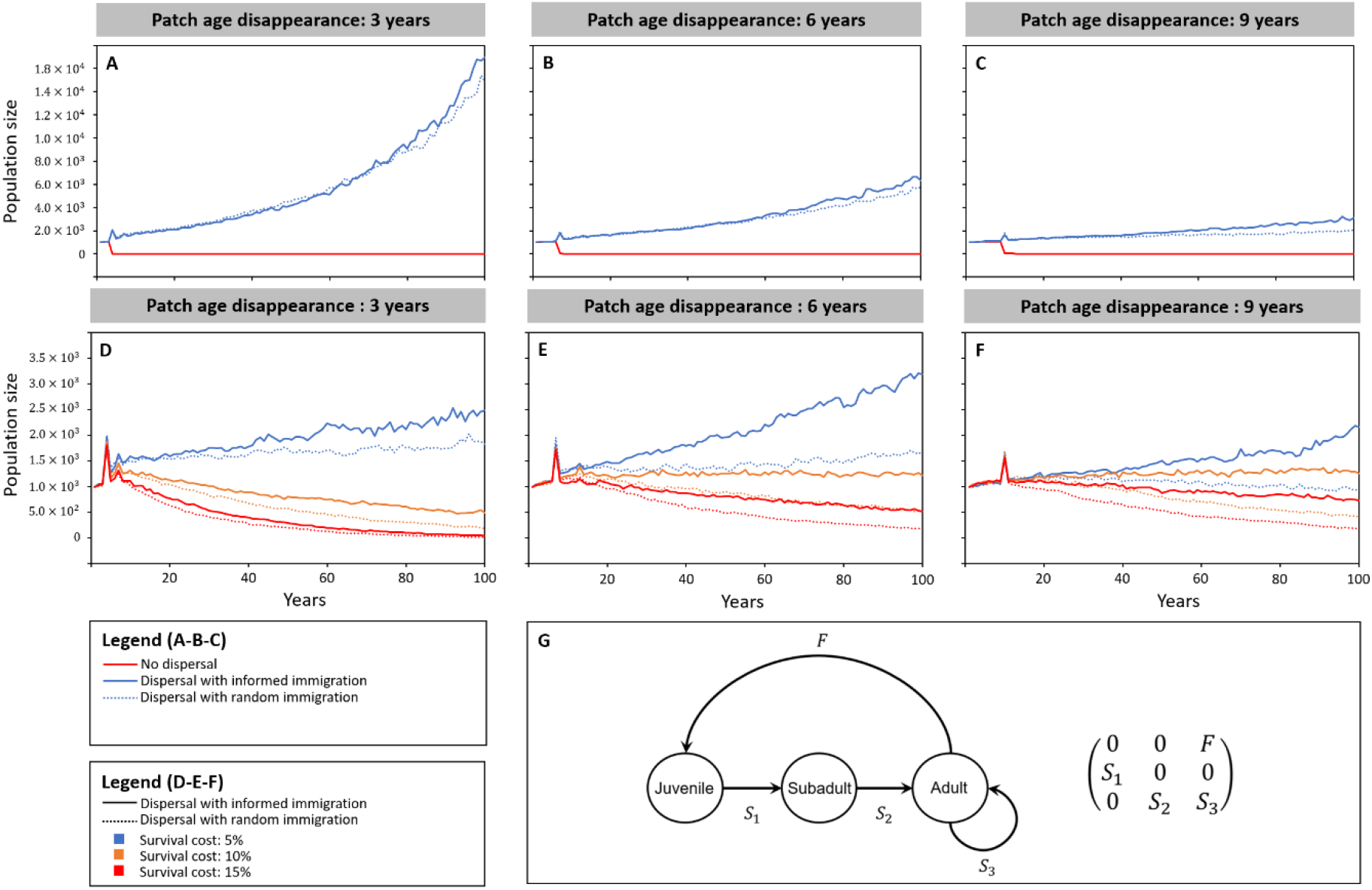
Effect of patch turnover on SSP size (number of adults) based on simulations. We considered three turnover rates: high (patch age before disappearance: 3 years, A and D), medium (6 years, B and E), and low (9 years, C and F). Three dispersal scenarios were also considered: (1) no dispersal (individuals were not able to escape when a patch disappeared and thus died); (2) dispersal with random immigration (individuals could disperse to escape the disappearance of a patch or disperse by choice even if the patch remained available. Immigration was random between the patches in the metapopulation and was not influenced by the age of the patch); (3) dispersal with informed immigration (similar to scenario 2, but individuals preferentially immigrated to recently created patches where fecundity F was highest). In scenarios 2 and 3, we considered two possible options regarding dispersal cost: (i) individuals did not incur any survival loss during dispersal (i.e. ‘non-costly dispersal’, A, B, and C); (ii) survival loss related to dispersal was either low (5% survival cost), medium (10%), or high (15%) (i.e. ‘costly dispersal’, D, E, and F). To investigate the demographic consequences of these scenarios, we used a female-dominant Leslie matrix (G) based on four demographic parameters: F = the female achieved fecundity, S_1_ = juvenile survival, S_2_ = subadult survival, and S_3_ = adult survival with both environmental and demographic stochasticity.

Our simulations also showed that an SSP’s extinction risk increased with the survival cost related to dispersal and was mitigated by ‘informed immigration’ (**Fig.4D, 4E**, and **4F**). In the scenarios with a high patch turnover rate (disappearance of patch after 3 years of availability), SSPs inevitably went extinct when the survival loss was higher than 10%. Random immigration accelerated the SSP’s decline compared to informed immigration directed toward recently created patches (where female fecundity was highest). In the scenarios with medium patch turnover (disappearance after 6 years of availability), the SSP decreased when the survival loss was 15% with both random and informed immigration. With a 10% survival loss, informed dispersal mitigated the decline of an SSP, whereas random dispersal drove the SSP to extinction. In the scenarios with low patch turnover (disappearance after 9 years of availability), the SSP experienced a marked decline only when immigration was random and the survival loss was equal to or higher than 10%. In summary, the simulations showed that patch turnover may result in SSP decline when the survival cost of dispersal is relatively high (10–15%) and immigration decisions are not adjusted to patch age and related fitness prospects.

### Phenotypic specialization in riverine and logging environments

We focused on three morphological characteristics known to condition movement capacity in amphibians: body size, body condition and relative leg size (Gomes et al. 2009, Hillman et al. 2014). Of the 400 tadpoles, 295 survived until metamorphosis, resulting in a relatively high metamorphosis success rate (*mean* ± *sd* = 0.75 ± 0.08), which did not differ according to the landscape type of origin (n = 400, *F*_1,3.8_ = 0, *P* = 0.95). Neither body size nor body condition at metamorphosis varied according to the landscape type (n = 295; body size: *F*_1,4_ = 0, *P* = 0.99; body condition: *F*_1,3.3_ = 1.12, *P* = 0.36). However, the allometric relationship between leg size and body size varied according to landscape type (n = 295; body size: *F*_1,227_ = 776.91, P < 0.0001; landscape type: *F*_1,3.4_ = 3.74, P = 0.13; body size × landscape type: *F*_1,227_ = 6.43, P = 0.012, **Fig. 5**). The larger the toadlets, the more those originating from logging SSPs tended to have longer hind limbs than those originating from riverine SSPs. Yet this difference was only just significant for large animals, as indicated by sliced tests (respectively performed at the 1^st^, 2^nd^ and 3^rd^ quartile of the size distribution: *F*_1,3.8_ = 1.64, P = 0.27; *F*_1,3.4_ = 3.28, P=0.15; *F*_1,4.01_ = 6.54; P = 0.06).

**Fig.5.**
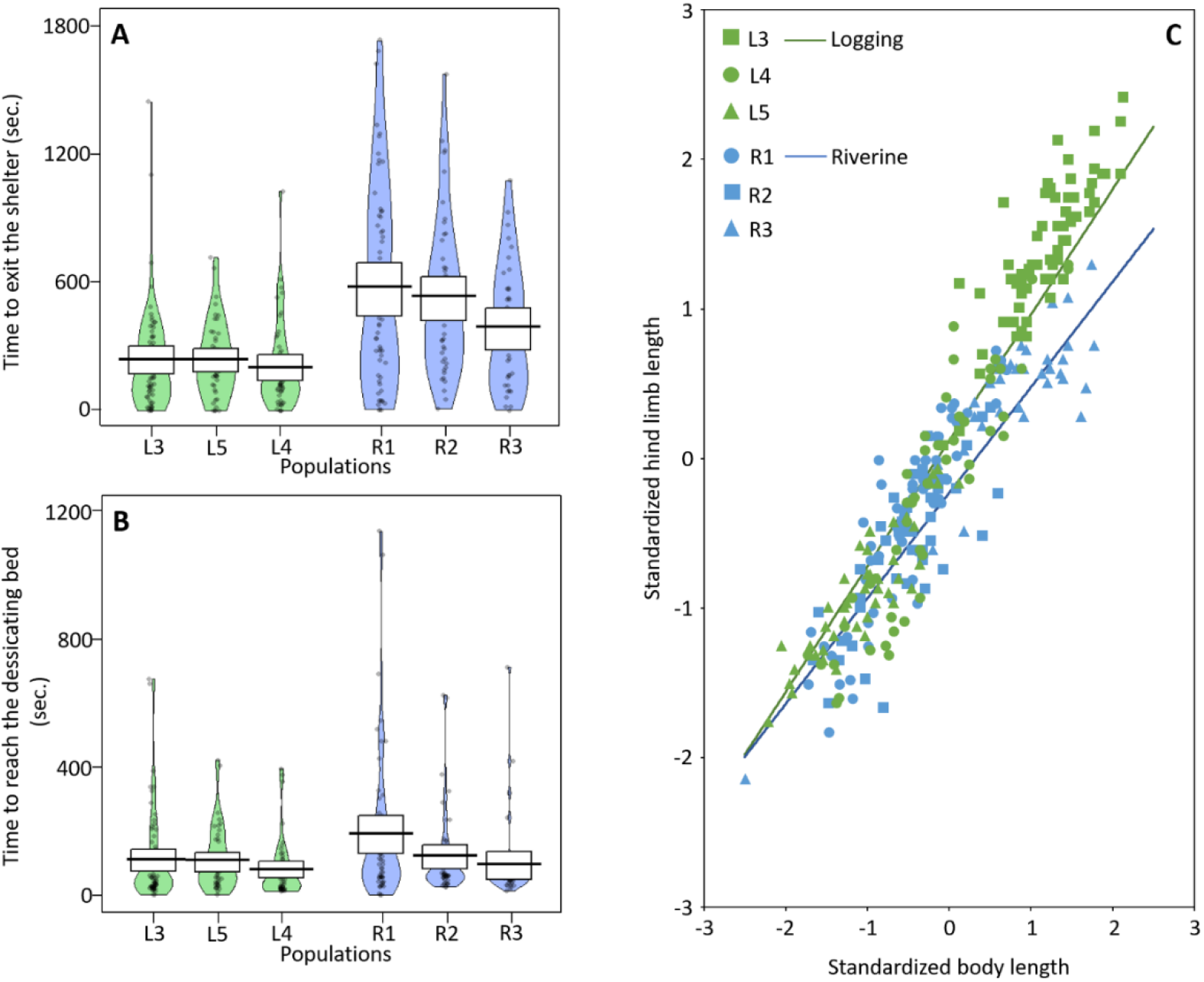
Behavioral and morphological specialization in six Bombina variegata SSPs occurring in logging environments (L3, L4, and L5) and riverine environments (R1, R2, and R3). We examined how the type of environment affected two behavioral traits: the time to exit the refuge chamber (A), a proxy for neophobia, and the time to reach the desiccation zone (B), a proxy for exploration propensity. We also examined how environment type affected hind limb length while considering body length as a control covariate in the model. These observations were recorded for 295 toadlets at metamorphosis.

Based on the behavioral trials in the experimental arenas, we measured variables along the boldness–shyness behavioral axis, a personality trait consistently involved in dispersal syndromes across different organisms (Cote et al. 2010). Toadlet behavior was characterized using three measures: one to assess neophobia (BEHAV1 = the time taken to leave the familiar refuge chamber) and the other two to assess exploration propensity (BEHAV2 = the time taken to reach the desiccant zone after leaving the refuge chamber; BEHAV3 = the time taken to cross the desiccant zone and get out of the arena). The findings showed that the neophobia of toadlets significantly varied according to their landscape of origin (**Fig.5**), but not according to their body size or its interactive effect with landscape (N=295; landscape type: F_1,3.5_=47.30, P=0.004; body size: F_1,13.9_=0.03, P=0.86; their interactive effect: F_1,12.1_=0.01, P=0.92). Individuals from riverine landscapes were 2.13+-0.24 times slower to leave the refuge chamber than those from logging landscapes (respectively 8.58+-0.67 mn and 3.89+-0.33 mn, see **Fig.5**). Once the toadlet left the refuge chamber, the latency time to reach the desiccating zone was very short (mean 1.83 +- 0.21 mn) and did not vary depending on the landscape of origin, body size, or their interactive effect (N=295; landscape type: F_1,3.9_=1.84, P=0.25; body size: F_1,30_=0.42, P=0.52; their interactive effect F_1,33.4_=0.26, P=0.61, see **Fig.5**). In contrast, the latency time to get out of the arena significantly varied depending on the landscape of origin, but not according to body size (N=295; landscape type: χ^2^_1df_=9.56, P =.002; body size: χ^2^_1df_=0.01, P=0.91; their interactive effect: χ^2^_1df_=0.28, P=0.59). The time taken – and thus the increased hazard – to get out of the arena for toadlets originating from logging SSPs was 1.93 times greater than for toadlets from riverine SSPs.

### Neutral genetic variation in riverine and logging environments

We investigated the neutral genetic footprint associated with each landscape type using microsatellite data collected in four SSPs: two from each landscape type (a total of 667 toads genotyped within L3 and L4 logging environments and R1 and R2 riverine environments; **Table S2**). Our analyses revealed, first, that riverine systems exhibited a higher deviation from the Hardy–Weinberg equilibrium than logging systems. Significant deviation from this equilibrium was detected for 75% and 45% of the loci in riverine SSPs R1 and R2 respectively, and 0% and 18% in the logging SSPs L3 and L4 (**Table S3**). Second, we found lower genetic diversity in riverine SSPs than in logging SSPs: both the allelic richness and the expected heterozygosity (uHE) were substantially lower in riverine SSPs. In contrast, inbreeding coefficients (*F_IS_*) were higher in riverine SSPs: 87% of the loci were found to have a lower uHE in riverine SSPs compared to logging SSPs (**Table S3**). Similarly, our parentage analyses also showed that individual inbreeding (LR test, *χ*^2^ = 32.55, P <0.0001; **Fig.6**) and the relatedness level within patches (LR test, *χ*^2^ = 43.14, P <0.0001, **Fig.7**) were drastically higher in riverine SSPs. In addition, riverine SSPs had a much smaller effective population size than logging SSPs: the effective population size was 9.7 (95% CI 6.4–13.7) in R1 and 22.6 (95% 19.3–26.3) in R2, while it was 138.9 (95% 80.2–353.6) in L4 and 168.6 (95% 129.9–229.3) in L3.

**Fig.6.**
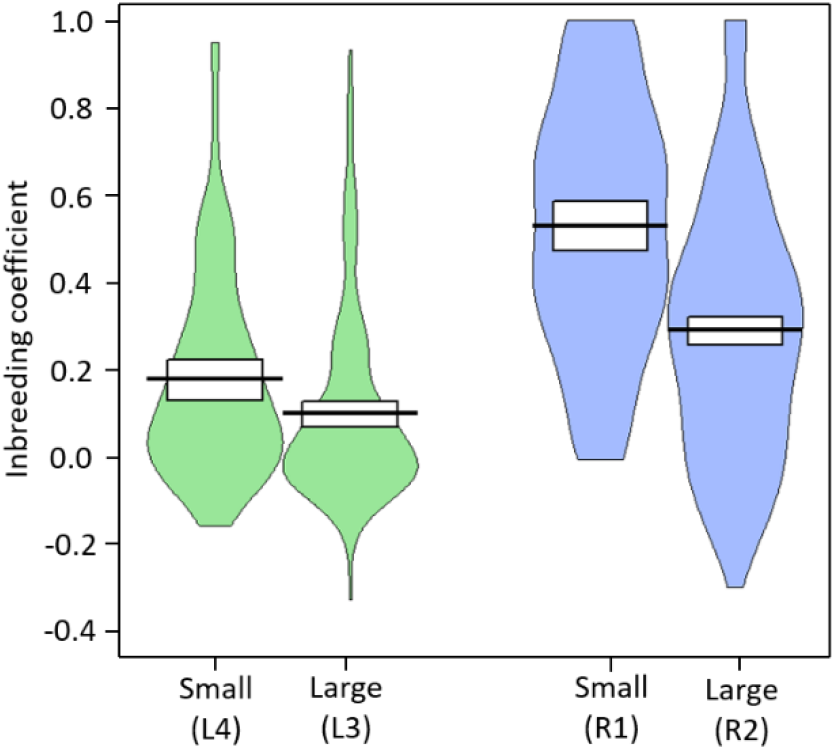
Inbreeding in the riverine (blue) and logging (green) *Bombina variegata* SSPs.

**Fig.7.**
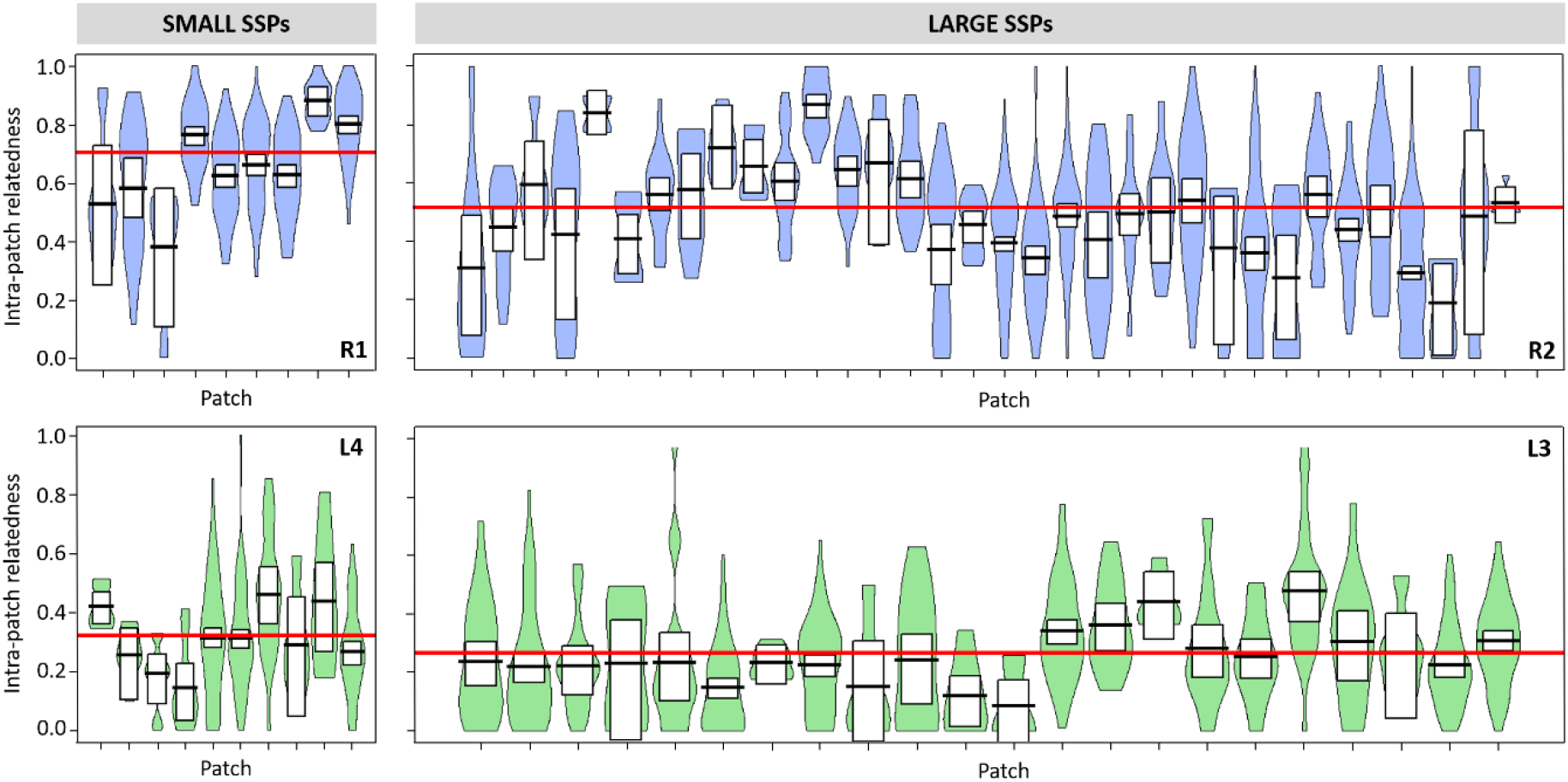
Relatedness within patches in the riverine (blue) and logging (green) *Bombina variegata* SSPs. Patches where less than 6 relatedness values were available were removed from the plot.

Within both types of environment, the size of SSPs (large: R2 and L3; small: R1 and L4) was also an important predictor of individual relatedness and inbreeding. The relatedness level within patches was higher (**Fig.7**) in small SSPs than in large SSPs (LR test, *χ^2^* = 214.38, P <0.0001), and this difference was larger in riverine SSPs (LR test, *χ*^2^ = 6.16, P = 0.01). Similarly, inbreeding within patches was higher (**Fig.6**) in small SSPs than in large SSPs (LR test, *χ*^2^ = 76.56, P <0.0001), and the interaction between these two factors was also confirmed (LR test, *χ*^2^ = 8.88, P = 0.01).

We then investigated how patterns of genetic differentiation between patches differed between logging and riverine SSPs. First, we used the Bayesian genetic clustering approach to examine hierarchical genetic structure in the SSPs. In the two SSPs from logging environments, we failed to detect any hierarchical genetic structure (K = 1) (**Fig.8**). In contrast, our analyses revealed a strong hierarchical genetic structure overlaying the spatial distribution of patches in the two SSPs from riverine environments (**Fig.8**). Second, we sought to verify that differences in spatial genetic patterns between landscapes were still significant while controlling for the potential difference of functional connectivity between SSPs using direct gradient analyses. These revealed that genetic differentiation correlated with geographic resistance between individuals whatever the landscape type, indicating substantial genetic isolation by distance even in logging systems (**Fig.9**). There was considerable support for the effect of the hydrological network on genetic differentiation in the large SSPs of both landscape types (SSPs R2 and L3), but not in the small SSPs whatever the landscape type (**Table 5**). Most importantly, even after correcting for these potential landscape effects, spatial genetic differentiation remained higher in riverine than in logging SSPs as revealed by the model averaged slope estimate associated with the geographic resistance effect (*mean* ± *sd*: 2.38 ± 0.07 for R1 and 2.55 ±0.03 for R2; 1.48 ± 0.15 for L3 and 0.68 ± 0.04 for L4).

**Fig.8.**
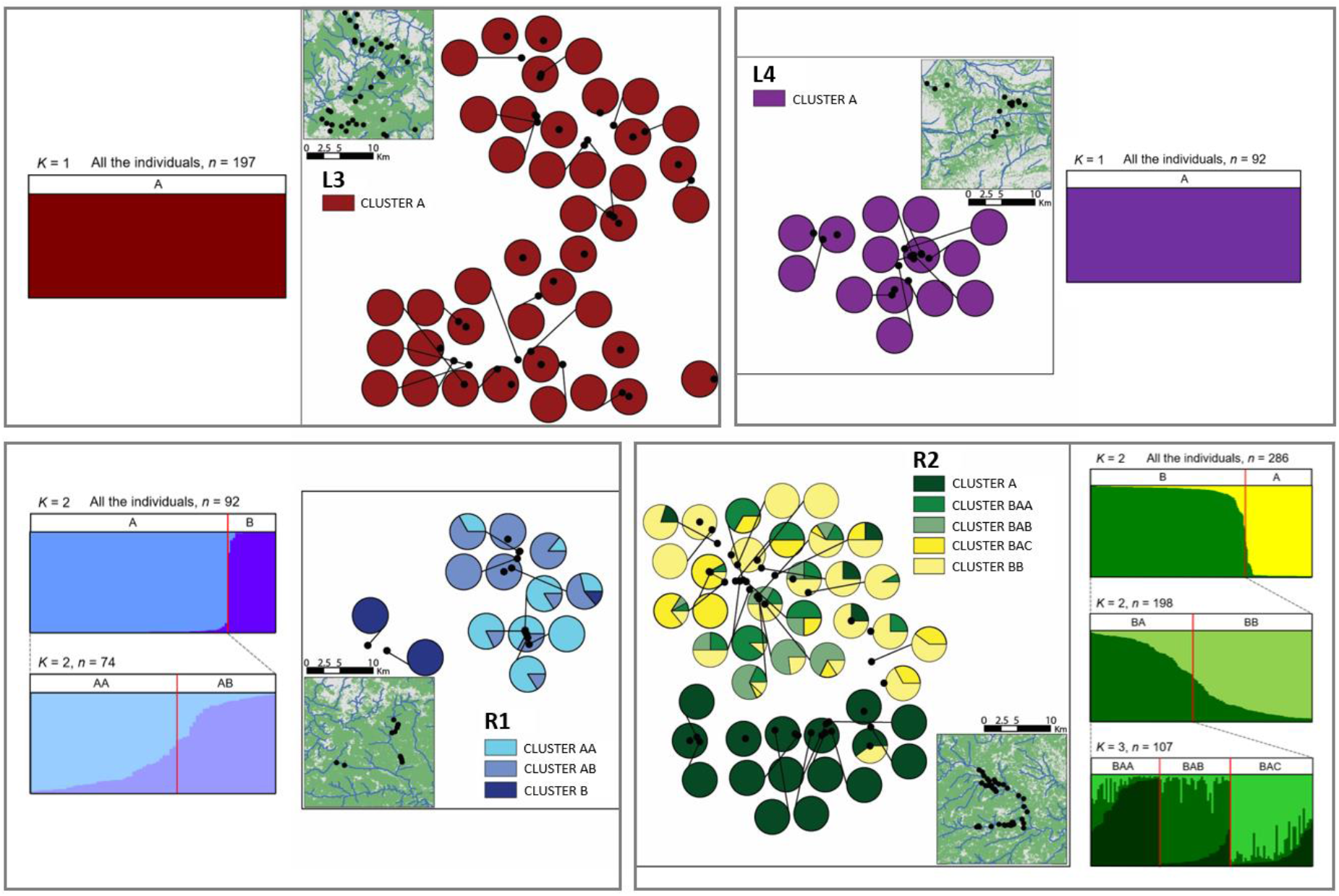
Patterns of neutral genetic variation and spatial distribution of the hierarchical genetic clusters in two riverine (R1 and R2) and logging SSPs (L3 and L4). The analyses were conducted using the program STRUCTURE. In L3 and L4, no genetic structuring was detected. In contrast, a complex genetic structure was found in R1 and R2. In R1, the hierarchical analysis revealed the existence of two initial genetic clusters (A and B), and genetic substructures within cluster A where two nested clusters were inferred (AA and AB). In R2, the hierarchical analysis highlighted the presence of two initial genetic clusters (A and B), and a substructure was further detected within cluster B (BA and BB). Genetic substructures were then identified within the cluster BA, with three additional genetic clusters inferred (BAA, BAB, and BAC).

**Fig.9.**
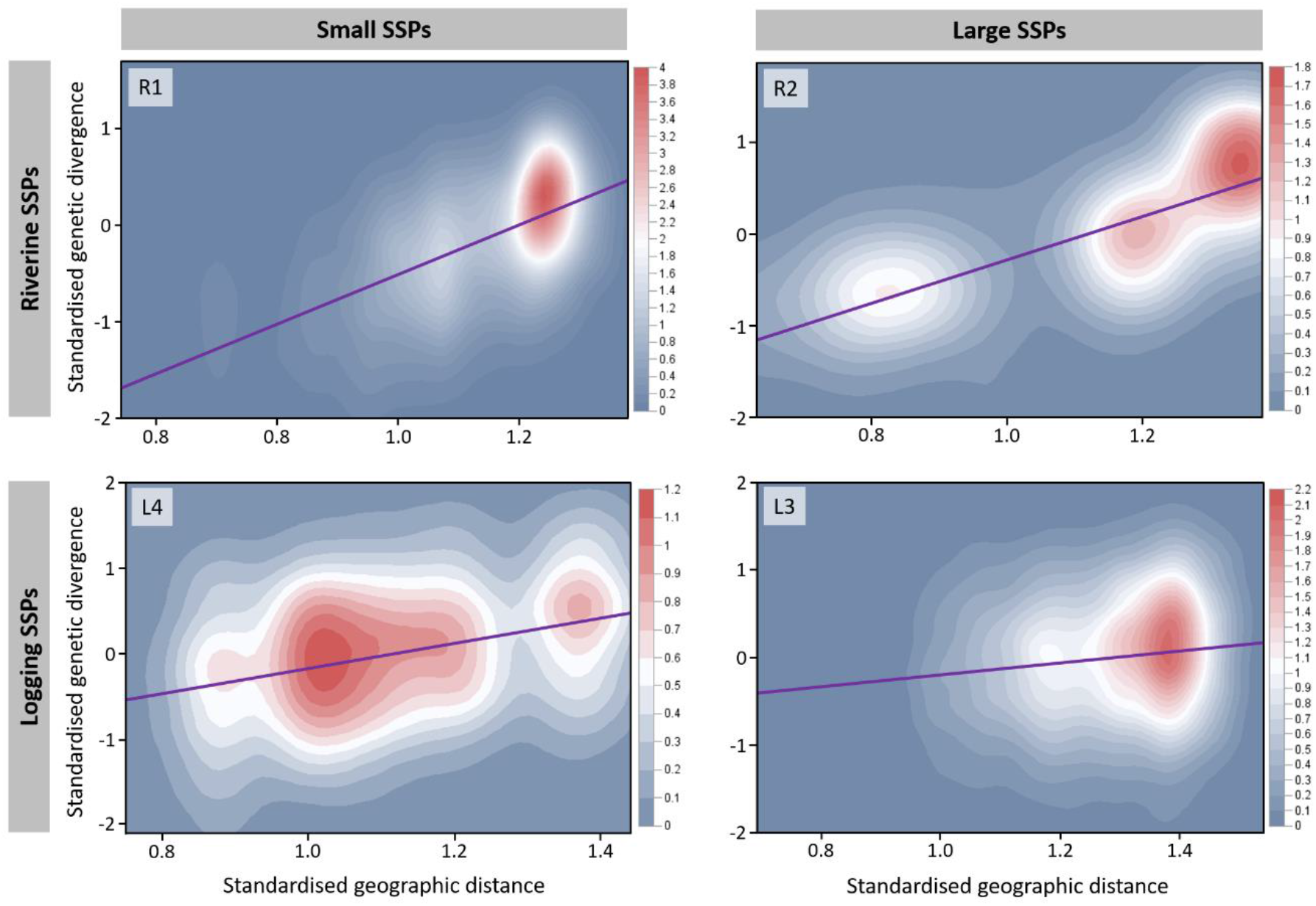
Genetic divergence according to geographic distance in two riverine (R1 and R2) and two logging SSPs (L3 and L4). Each figure represents the contour plot of the kernel density bivariate estimates between the pairwise genetic distance and the pairwise geographic resistance for each SSP. Kernel densities were estimated using a Gaussian distribution. The graduated color contour indicating the (smoothed) observation count is presented on the right side of each plot. The line represents the predicted regression curve between the genetic distance and the geographic resistance from the linear mixed model estimates.

**Table 5.** Relative support of the mixed models for each of the four SSPs and their related estimates of geographic resistance (L3 and L4 = logging SSPs, R1 and R2 = riverine SSPs, see Map S1 for details). ‘Model’ indicates the effect introduced in the model in addition to geographic resistance (null = the model including only the effect of geographic resistance). ‘Waic’ indicates the weighted AIC of the model. 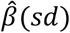 indicates the beta weight associated with the effect of geographic resistance and its standard deviation. The variable ‘slope ‘ corresponds to the effect of both geographic distance and topographic roughness. The variable ‘network’ corresponds to the effect of the hydrological network.

| Model | Waic | $\hat{\beta}(sd)$ |
| --- | --- | --- |
| **SSP L3** |  |  |
| null | 0.469 | 1.4763 (0.1544) |
| slope | 0.245 | 1.4811 (0.1545) |
| network | 0.191 | 1.4798 (0.1545) |
| slope+network | 0.095 | 1.4826 (0.1546) |
| **SSP L4** |  |  |
| null | 0.169 | 0.6783 (0.0415) |
| slope | 0.084 | 0.679 (0.0415) |
| network | 0.507 | 0.6764 (0.0415) |
| slope+network | 0.240 | 0.677 (0.0415) |
| **SSP R1** |  |  |
| null | 0.391 | 2.381 (0.0713) |
| slope | 0.276 | 2.386 (0.0714) |
| network | 0.226 | 2.3845 (0.0714) |
| slope+network | 0.107 | 2.3862 (0.0714) |
| **SSP R2** |  |  |
| null | 0.070 | 2.5459 (0.0251) |
| slope | 0.030 | 2.5461 (0.0251) |
| network | 0.601 | 2.5479 (0.0251) |
| slope+network | 0.299 | 2.5482 (0.0251) |

## Discussion

Taken together, the results of this study provide for the first time an extended picture of the effect of anthropogenic disturbance on dispersal in a vertebrate, from dispersal-related phenotypic specialization expressed early in life, through the dispersal pattern emerging in spatially structured populations, to the genetic footprint arising throughout successive generations. Overall, this analysis revealed that anthropogenic disturbance not only strongly promotes dispersal throughout a toad’s lifetime, but also prenatally enhances a toadlet’s risk-proneness and, to a certain extent, favors longer hind limb length at metamorphosis. Another finding was that gene flow also substantially increased in anthropogenic landscapes independently of the SSP’s size or functional connectivity.

### Natal and breeding dispersal rates and distances in SSPs depend on patch turnover rate

The results revealed contrasting dispersal patterns between riverine and logging SSPs. In the demographic component of our study, the sampling protocol was weakened by a potential confounding effect of the population’s position along the latitudinal gradient with its status (logging/riverine). However, we can rule out the possibility of an effect of latitude on the dispersal patterns drawn in our analyses, as the molecular inferences showed drastically increased gene flow (resulting from dispersal) in an SSP at a low latitude (L4), which indicates that latitude has a marginal effect on effective dispersal within SSPs. We are therefore confident in the reliability of our results regarding the effect of turnover rate on dispersal patterns.

In riverine SSPs, we observed a complete suppression of natal dispersal, as well as very low breeding dispersal, together with a reduced dispersal kernel – the opposite of the dispersal pattern observed in logging SSPs. In the latter, we nevertheless found some substantial inter-population variation regarding the contribution of natal dispersal to the overall dispersal process and to seasonal variation in dispersal rates. Such differences between logging SSPs likely reflect variation in the anthropogenic disturbance regime resulting from local woodland management practices. Indeed, patch turnover depends on both the extent and the frequency of the logging activities that create patches, as well as the post-logging rehabilitation operations that may lead to patch destruction (e.g. filling in of ruts used as temporary breeding ponds) (Cayuela et al 2018b). This likely results in a continuum of dispersal strategies along a gradient of patch disturbance, ranging from the near suppression of dispersal in riverine SSPs to a very high dispersal rate and long dispersal distances in some logging SSPs (e.g. L1 SSP). This pattern is thus very similar to that observed for wing dimorphism in insects alongside patch temporality gradients (Denno et al. 1996, Roff & Fairban 2007), in which dispersal is suppressed (i.e. high rate of wingless forms) in persistent habitats, while it is enhanced (i.e. high rate of winged forms) in more ephemeral habitats according to the (natural) disturbance regime experienced at the landscape level.

### Phenotypic specialization in SSPs depends on patch turnover rate

Previous studies on *Bombina variegata* (Cayuela et al. 2018b) have revealed that dispersal events in logging SSPs are a mixture of departures conditioned by patch disappearance and unconditional departures that occur well before patch disappearance. The findings from our common garden experiments highlight that patch turnover rate prenatally mediates dispersal-related phenotypes and leads to phenotypic parallelism in logging and riverine SSPs. In particular, toadlets originating from logging SSPs exhibited higher risk-proneness than those from riverine SSPs, as revealed by their swiftness in leaving the refuge chamber and in getting out of the arena after crossing a harsh substrate. Either neophilia or boldness (see Kelleher et al. 2018 for personality traits in amphibians) could explain the elevated risk-proneness we observed in logging SSPs. Disentangling these two personality traits is not straightforward (Greggor et al. 2015, Yuen et al. 2017), and further investigations would be useful to address this. Regardless of the exact composition of personality traits behind our identification of risk-prone behavior, our results clearly indicate behavioral specialization early in life according to the disturbance regime prevailing in the landscape.

Concerning morphological traits, we did not find any differences in body size and condition of toadlets from logging and riverine SSPs. However, the findings showed that toadlets innately have longer hind limbs in logging than in riverine SSPs. In anurans, hind-limb length is usually positively associated with locomotor performance (Choi et al. 2003, Philips et al. 2006, Gomes et al 2009, Hudson et al. 2016), and has also been found to be subject to rapid evolution at the edge of the invasion front in the introduced species *Rhinella marina* (Philips et al. 2006). Our results thus suggest that long hind limbs could be a phenotypic trait facilitating dispersal in logging habitats. Yet this difference in leg length between toadlets from the two environments was only observed in large individuals and was subject to substantial variation between SSPs. This significant but weak effect of anthropogenic disturbance on leg length could result from developmental constraints. Limb size mainly depends on the duration of the larval period in anurans, so species specialized for ephemeral pools (such as *Bombina variegata*), which are selected for fast larval development, usually exhibit shorter hind limbs compared to other species (Gomez-Mestre & Buchholz 2006).

Overall, dispersal and related behavioral traits are usually highly plastic phenotypes subject to partial genetic control (reviewed in Saastamoinen et al. 2018), i.e. determined by G × E interactions. Therefore, the phenotypic specialization highlighted in our study may have a genetic basis and/or may be associated with transgenerational plasticity. In the absence of crossbreed design, we were not able to disentangle the relative contribution of maternal effect and parental genotypes in the phenotypic variation observed in logging and riverine SSPs. However, it is possible to rule out the hypothesis of transgenerational plasticity mediated by toadlet body size at metamorphosis. In amphibians, female energy investment in breeding influences the size of eggs and the amount of energetic resources available for the development of embryos and larvae before they become fully heterotrophic (Kaplan 1987, 1992). Studies have reported a positive relationship between egg size and offspring body size at metamorphosis due to carry-over effects (Laugen et al. 2005, Räsänen et al. 2005, Dziminski et al. 2006), and body size is an important predictor of behavioral traits (e.g. exploration propensity and risk-taking behavior) related to dispersal (Kelleher et al. 2017, 2018). In our case study, no difference in body size was detected in toadlets from logging and riverine environments, which indicates that patch turnover rate is not likely to alter individual behavior due to the morphological state at metamorphosis in SSPs. However, epigenetic factors (e.g. DNA methylation, micro-ARN, and histone structure) independent from female energy investment strategies may lead to transgenerational dispersal plasticity and contribute to the expression of phenotypic traits that facilitate or hinder dispersal over generations (Saastamoinen et al. 2018, Cayuela et al. 2019a). Yet it is very likely that a genetic basis partially determines phenotypic specialization in logging and riverine environments, resulting in incomplete genetic parallelism between SSPs (Elmer & Meyer 2011, Conte et al. 2012). First, transgenerational plasticity in dispersal-related traits is usually subject to genetic control (Cayuela et al. 2019a), likely due to a strong association between epigenetic variation and genetic variants in *cis* and *trans* (Dubin et al. 2015, Zaghlool et al. 2016). Second, as predicted by the dispersal theory and reported in other study systems, dispersal is partly genetically determined, and behavioral traits related to dispersal often have a polygenic basis (Saastamoinen et al. 2018). Future studies using Next-Generation Sequencing approaches (Morozova & Marra 2008, Metzker 2010) could be undertaken to determine the role of genetic, transcriptional, and epigenetic variation in the disturbance-dependent phenotypic changes and dispersal evolution in *B. variegata*.

### Patch turnover and related dispersal costs and benefits determine SSP dynamics and persistence

Our simulations showed that the absence of dispersal inevitably leads to SSP extinction, and that the extinction speed increases with patch turnover. When dispersal was possible and had no survival cost, patch turnover had a positive effect on SSP size. This resulted from an increase in average fecundity due to the positive relationship between fecundity and patch age reported in logging SSPs (Boualit et al. 2019) and considered in our models. These results are congruent with field observations reporting that *Bombina variegata* SSPs may be very large (thousands of adults) in harvested woodlands in western Europe.

The models also showed that the risk of SSP extinction increased with the survival cost related to dispersal and was mitigated by informed immigration. This result is congruent with theoretical models and empirical evidence showing that dispersal only evolves if the benefits of moving outweigh the related costs (Bonte et al. 2012). In logging SSPs, dispersal seems to be favored as its costs are likely offset by the benefits of colonizing new patches. Survival (at juvenile, subadult, and adult stage) is lower in logging than in riverine SSPs (Cayuela et al. 2016a), which likely results from mortality caused by logging operations and dispersal-related mortality (Cayuela et al. 2018b). This survival cost of dispersal can be direct, i.e. associated with movement in the landscape matrix (Bonte et al. 2012). It can be also indirect, resulting from the high energy allocation necessary for recurrent dispersal events over the toad’s lifespan, which might lead to earlier and stronger senescence due to trade-offs (Cayuela et al. 2019b). These direct and indirect costs are obviously offset when animals are forced to disperse subsequently to patch destruction resulting from logging practices (Cayuela et al 2018b). It should be noted that even when patches remain available, dispersal may still be strongly favored since their suitability rapidly declines over time due to the natural silting of ruts if these are not regularly disturbed by vehicle traffic (Boualit et al. 2019). Furthermore, even if patch suitability is sustained by regular human disturbance (Boualit et al. 2019), dispersal costs might be mitigated through the colonisation of recently created patches in which fitness prospects are likely enhanced due to density-dependent mechanisms. A low density of adults in newly available patches likely reduces the risk of larval competition, which is an important driver of metamorphosis success (Jasieński 1988) – for this reason adults preferentially reproduce in tadpole-free waterbodies (Cayuela et al. 2016d, 2017b).

These potential benefits likely favor the evolution of dispersal and dispersal-enhancing morphological and behavioral traits (i.e. ‘dispersal syndromes’) in logging SSPs. They also likely contribute to context-dependent dispersal (Clobert et al. 2009) and matching habitat choice (Edelaar et al. 2008), implying that individuals adjust their dispersal decisions according to the local fitness prospects determined by the biotic and abiotic characteristics of breeding patches. This prediction has been verified by two studies reporting context-dependent dispersal in logging SSPs (Tournier et al. 2017, Boualit et al. 2019). In particular, these studies showed that adult emigration and immigration decisions depend on a pond’s hydroperiod and the size and annual disturbance of the patch: three factors that locally affect juvenile recruitment and very likely individual fitness.

In riverine SSPs, the near absence of both natal and breeding dispersal suggests that the benefits of dispersal do not compensate for its potential costs. First, the absence of patch loss resulting from anthropogenic or natural processes means individuals are not forced to disperse to survive and reproduce. In river environments, the local fitness prospects do not deteriorate with patch age, as the process of natural silting of rocky pools is frequently interrupted by river flooding occurring outside the breeding period (Cayuela et al. 2011, 2015a). This makes pools available for breeding from one year to another and limits the risk of larval mortality caused by desiccation. It is also possible that the near absence of dispersal results from variability in pond characteristics (e.g. hydroperiod and temperature; Cayuela et al. 2011) within a patch. Indeed, the benefit of dispersing would be low if the variability in environmental conditions would be similar at intra-patch and inter-patch scales. Overall, the lack of apparent compensatory benefits should not favor and may even counter-select for dispersal and dispersal-enhancing traits in riverine SSPs. This is in line with our results, which show that riverine toadlets innately display low risk-taking behavior and have short hind limbs, two phenotypic traits hindering dispersal in amphibians (Cayuela et al. 2018c).

### Genetic variation patterns in SSPs depend on patch turnover rate

Our study showed for the first time that the spatiotemporal dynamics of habitat patches in a landscape may have an effect at least as important as landscape fragmentation on gene flow patterns. The findings highlighted that low dispersal rates and distances are associated with a weaker genetic structure and lower IBD in riverine than in logging SSPs. Our sampling design allowed us to disentangle the relative effects of gene flow and genetic drift, which both contribute to genetic differentiation within SSPs (Slatkin 1977, Broquet & Petit 2009). As well as genetic structure differences caused by patch turnover, we found higher IBD in the two small SSPs (L4 and R1) than in the two large SSPs in each environment.

Our analysis also took into account the functional connectivity within SSPs by considering two landscape factors (i.e. topography and hydrological network) that are critically important to the genetic structure of amphibian populations (reviewed in Cayuela et al. 2018c). As reported in 20 previous studies (Cayuela et al. 2018c), genetic differentiation increases with increasing topographic slope as this raises the energy cost of dispersal during the transience phase in amphibians. In addition, our results showed that genetic differentiation was better explained by the distance of the hydrological network than the Euclidean distance between two patches. This indicates that the hydrological network improves the functional connectivity between patches by reducing the cost of displacement in the landscape matrix, which is congruent with the findings of six previous studies (Cayuela et al. 2018c). Interestingly, genetic differentiation nevertheless remained higher in riverine than in logging SSPs, despite a denser hydrological network and therefore weaker landscape resistance in riverine SSPs. This suggests that in our study system, the rate of patch turnover may be a more important driver of neutral genetic variation than landscape connectivity.

Our findings suggest that variation in the turnover rate of SSPs has far-reaching consequences on the evolutionary forces involved in the migration–selection–drift balance. In riverine SSPs, reduced gene flow between patches leads to lower genetic diversity and smaller effective population size compared to logging SSPs. Such low standing genetic variation could limit the adaptive response of riverine SSPs to novel environmental conditions. The probability of allele fixation increases with the magnitude of the beneficial effect and the effective population size, and this probability is significantly higher when the allele has a high initial frequency (Barrett & Schluter 2008, Hedrick 2013). Moreover, standing genetic variation usually allows faster adaptation as beneficial alleles are already present in the population (Barrett & Schluter 2008, Hedrick 2013). Overall, our results suggest that riverine SSPs should suffer from a lower capacity of ‘evolutionary rescue’ (Carlson et al. 2014) than logging SSPs, which could increase their sensitivity to current global changes.

The results also revealed a high level of inbreeding and relatedness within riverine SSPs, confirming the results of a previous study (Cayuela et al. 2017a). These findings raise important questions about the mechanisms associated with the repression of inbreeding depression. As mentioned, the survival rate at all life stages (juvenile, subadult, and adult) is higher in riverine than in logging SSPs (Cayuela et al. 2016a), suggesting a marginal effect of inbreeding on postmetamorphic survival and negligible inbreeding depression. A previous study conducted in the R2 SSP indicated that the absence of disassortative mating does not seem to mitigate inbreeding risk: females even prefer to reproduce with related males from their own patch (Cayuela et al. 2017a). Although the effect of inbreeding on reproductive performance remains as yet unevaluated, those results suggest a low genetic load in riverine SSPs. This could be due to high efficiency in purging deleterious alleles and the genomic architecture of genetic load, especially a low linkage of deleterious recessive alleles (Bersabé et al. 2016, Hedrick & Garcia-Dorado 2016). Future studies could be carried out to identify the mechanisms involved in the genetic purging of inbreeding depression resulting from limited dispersal in riverine SSPs.

## Conclusion

Our study showed that anthropogenic habitat disturbance is likely an important driver of dispersal evolution. The results found that, like landscape fragmentation, human-driven variation in patch turnover may promote morphological and behavioral specialization related to dispersal. In particular, it may lead to phenotypic parallelism affecting dispersal syndromes and patterns (dispersal rate and distance) in SSPs exposed to contrasting levels of patch turnover. This phenotypic parallelism is likely underpinned by genetic and/or epigenetic parallelism, for which the molecular basis remains to be investigated. Our results also revealed that differences in dispersal patterns are associated with variation in the genetic structure of SSPs, which might affect local eco-evolutionary dynamics (Legrand et al. 2017). In particular, high gene flow and reduced effects of genetic drift allow higher genetic polymorphism to be maintained in SSPs experiencing high patch turnover than in SSPs with low patch turnover. In parallel, larger effective population size is expected to increase selection effectiveness in SSPs exposed to high patch turnover, giving them higher evolutionary potential and increased chances of evolutionary rescue in the case of environmental change (Carlson et al. 2014). These results emphasize the central role of anthropogenic disturbance in the spatiotemporal dynamics of landscapes and the related ecological and evolutionary processes.

In terms of the implications for conservation, the findings suggest that the approach may need to differ depending on the demographic and genetic characteristics of SSPs driven by the level of patch turnover. Those SSPs exposed to high patch turnover are likely to be highly sensitive to land use changes that result in a loss of functional connectivity between patches. An increase in landscape resistance reduces the success of dispersal following the disappearance of a patch and diminishes the chance of new patch colonization, which enhances the extinction risk of the whole SSP. In contrast, SSPs experiencing a low level of patch turnover suffer a loss of genetic diversity and high inbreeding, which may increase their sensitivity to environmental change (e.g. climate change or the emergence of new pathogens), potentially speeding up their extinction if they experience a demographic decline (i.e. inbreeding depression; Reed & Frankham 2003, Frankham 2010). This underlines the tremendous importance of adjusting conservation decisions to the demographic and genetic characteristics of SSPs adapted to contrasting regimes of habitat disturbance.

## Acknowledgments

We would like to thank all the undergraduate students and fieldworkers that contributed to the capture–recapture surveys, DNA samples, and common garden behavioral tests. This research project was funded by the Office National des Forêts, the Lorraine Direction Régionale de l’Environnement, de l’Aménagement et du Logement (DREAL), the Agence de l’Eau Rhin-Meuse, the Conseil Régional de Lorraine, the Conseil Régional de Champagne-Ardenne, the Conseil Régional de Picardie, the Conseil Général de l’Aisne, the Conseil Général d’Ardèche, the Conseil Général d’Isère and the Communauté de Communes de l’Argonne Ardennaise (2C2A). The study was authorized by three government environmental agencies: the Lorraine DREAL, the Rhône-Alpes DREAL and the Agence de l’Eau Rhin-Meuse. The ethical committee of Université Claude Bernard Lyon 1 approved the capture–recapture and common garden protocols.

