## Supplementary_material for "Anthropogenic disturbance drives dispersal syndromes, demography, and gene flow in spatially structured amphibian populations"

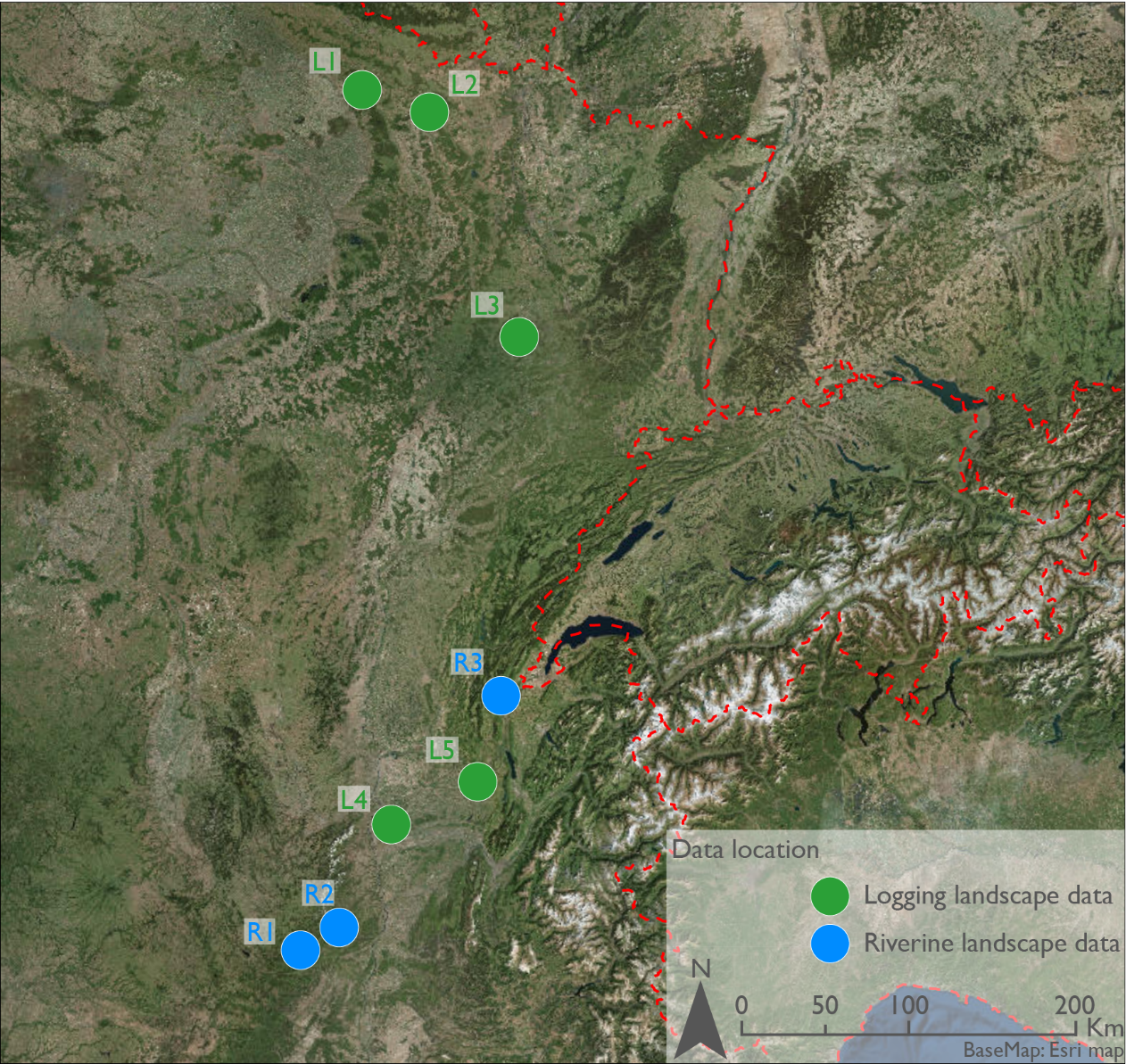

**Figure S1.** Map showing the eight studied SSPs in logging (L1 to L5) and riverine (R1 to R3) landscapes in France.

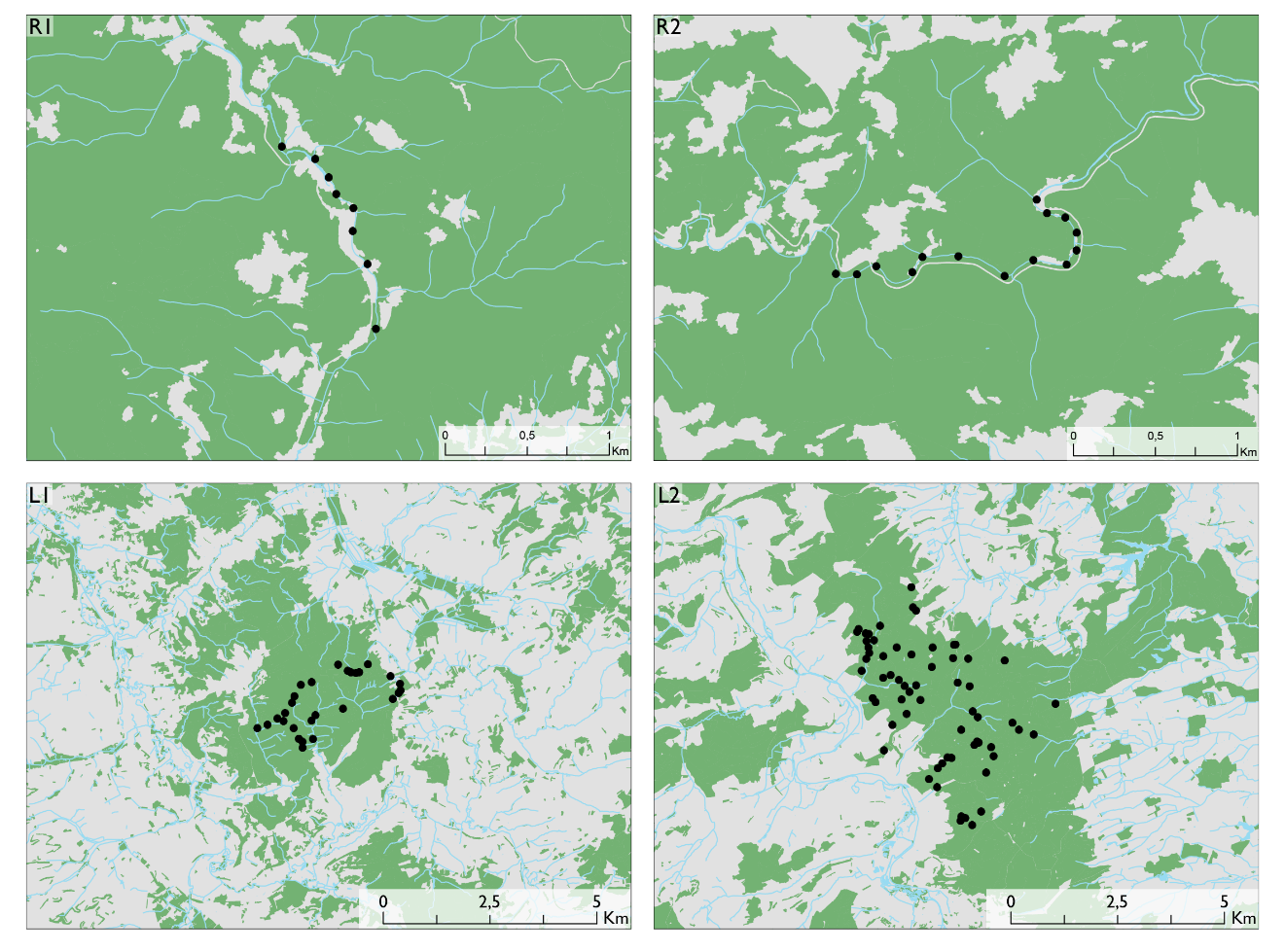

**Figure S2.** Map showing the four studied SSPs in logging (L1 to L2) and riverine (R1 to R2) landscapes surveyed using the capture–recapture method. The black spots correspond to breeding patches (i.e. groups of waterbodies). The green areas correspond to woodland.

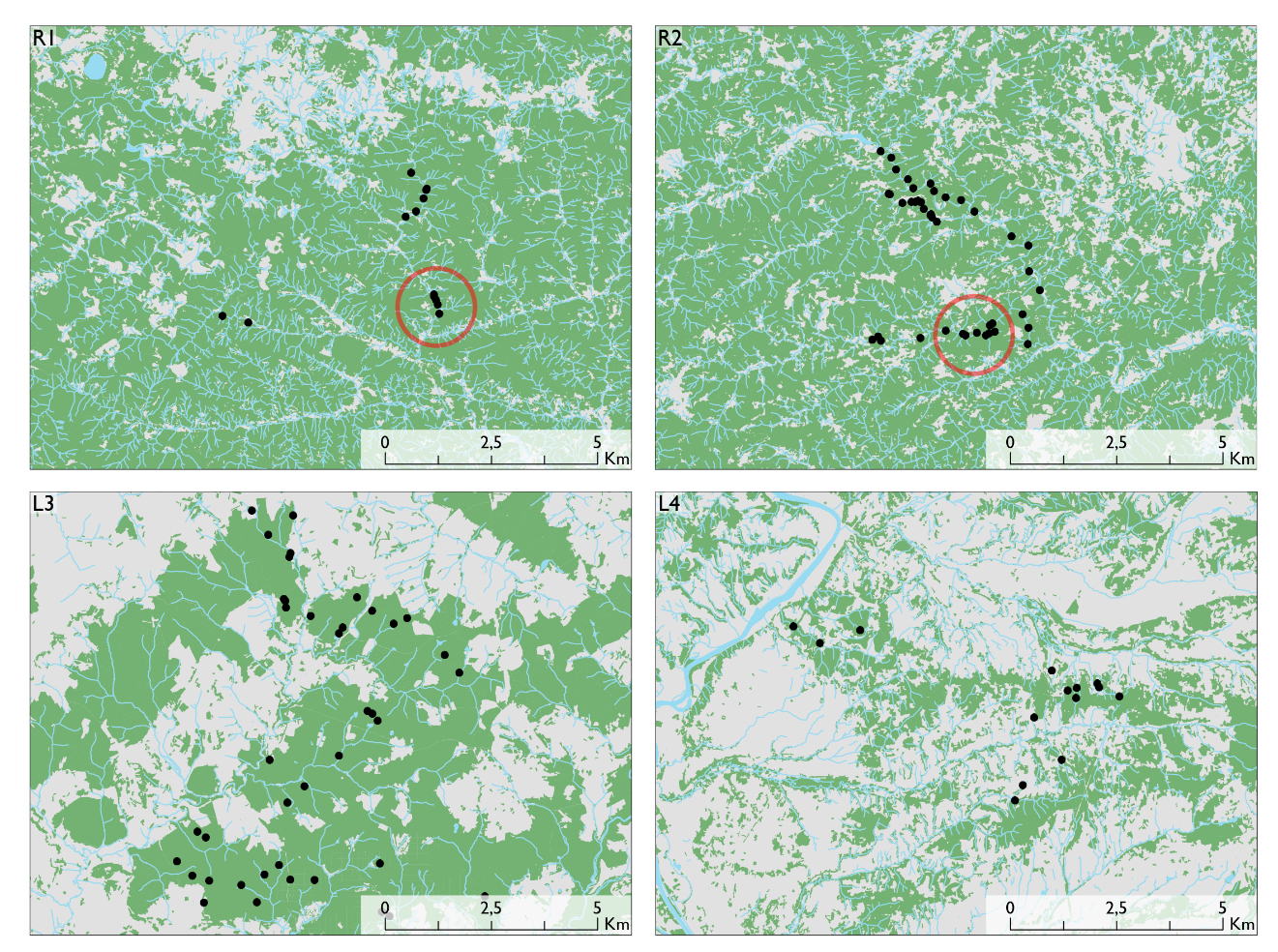

**Figure S3.** Map showing four studied SSPs in logging (L3 to L4) and riverine (R1 to R2) landscapes studied with genetic data. The black spots correspond to breeding patches (i.e. groups of waterbodies). The green areas correspond to woodland. In R1 and R2, the set of patches circled in red corresponds to patches that were surveyed using the capture–recapture method.

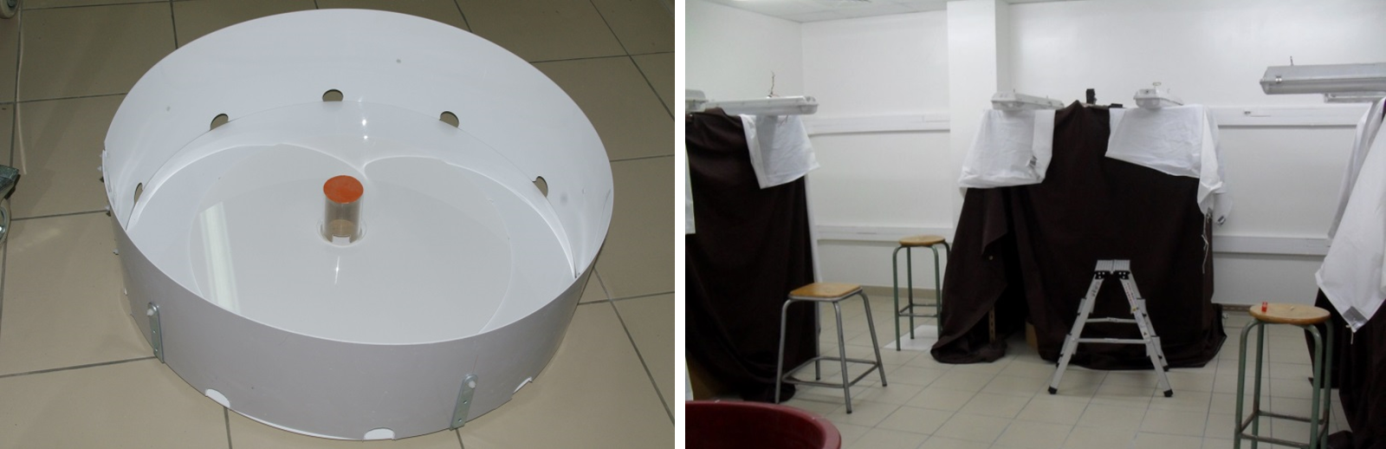

**Figure S4***.* The experimental arena for the behavioral assays (left) and the iron chamber in which the arena was enclosed (right).

**Table S1.** Number of juveniles, subadults and adults captured each year in the four SSPs (L1, L2, R1 and R2) considered in the study.

|  | Year 1 | Year 2 | Year 3 | Year 4 | Year 5 | Year 6 | Year 7 | Year 8 | Year 9 |
| --- | --- | --- | --- | --- | --- | --- | --- | --- | --- |
| L1 |  |  |  |  |  |  |  |  |  |
| Juv | 39 | 24 | 2 | 4 | 0 | 0 | 60 | 14 | 61 |
| Sub | 4 | 17 | 7 | 0 | 1 | 1 | 25 | 44 | 33 |
| Adu | 75 | 73 | 70 | 30 | 15 | 28 | 55 | 122 | 149 |
| L2 |  |  |  |  |  |  |  |  |  |
| Juv | 599 | 1006 | 945 | 295 | 40 | - | - | - | - |
| Sub | 114 | 534 | 827 | 775 | 511 | - | - | - | - |
| Adu | 1209 | 947 | 1177 | 1155 | 2194 | - | - | - | - |
| R1 |  |  |  |  |  |  |  |  |  |
| Juv | 237 | 86 | 417 | 82 | 12 | - | - | - | - |
| Sub | 99 | 192 | 110 | 187 | 53 | - | - | - | - |
| Adu | 764 | 647 | 789 | 480 | 592 | - | - | - | - |
| R2 |  |  |  |  |  |  |  |  |  |
| Juv | 278 | 47 | 138 | 132 | 46 | - | - | - | - |
| Sub | 19 | 381 | 65 | 85 | 76 | - | - | - | - |
| Adu | 596 | 573 | 512 | 455 | 581 | - | - | - | - |

**Table S2**. Number of patches and DNA samples in the four SSPs in riverine (R1 and R2) and logging environments (L3 and L4).

| SSP | Number of patches | Number of DNA samples |
| --- | --- | --- |
| R1 (small SSP) | 14 | 92 |
| R2 (large SSP) | 41 | 286 |
| L3 (small SSP) | 14 | 92 |
| L4 (large SSP) | 41 | 197 |

**Table S3**. Characteristics of the microsatellite markers developed in Cayuela et al. (2017) (the first eight from the left) and those published by Nürnberger et al. (2003) (Bv11.7, Bv11.2, Bv24.12) and Hauswaldt et al. (2007) (Bobom5F, Bobom8A, Bobom9H, Bobom10F, Bobom12F) in metapopulations experiencing unpredictable environments (R1 and R2) and predictable environments (L3 and L4). $N_{a}$ = mean number of alleles per locus, $N_{e}$ = effective number of alleles per locus, $H_{o}$ = observed heterozygosity, $H_{e}$ = expected heterozygosity, HWE = deviation from Hardy–Weinberg equilibrium (*** = significant, ns = non-significant), $F_{\mathrm{IS}}$ = inbreeding coefficient, Null = probability of null allele presence.

|  |  | Bomvar_DLWD0 | Bomvar_Cons470 | Bomvar_CUSGH | Bomvar_DORC3 | Bomvar_DM3QM | Bomvar_4EMX7J | Bomvar_CWL3Z | Bomvar_4EGYJ3 | bv11.7 | bv11.2 | Bv24.12 | Bobom8A | Bobom12F | Bobom10F | Bobom5F | Bobom9H |
| --- | --- | --- | --- | --- | --- | --- | --- | --- | --- | --- | --- | --- | --- | --- | --- | --- | --- |
| R1 | $N_{a}$ | 2 | 2 | 2 | 4 | 2 | 3 | 2 | 3 | 3 | 2 | 2 | 3 | 8 | 6 | 3 | 6 |
|  | $N_{e}$ | 1.49 | 1.63 | 1.18 | 1.57 | 1.65 | 1.02 | 1.13 | 1.07 | 1.35 | 1.24 | 1.86 | 1.43 | 1.77 | 2.39 | 1.23 | 1.95 |
|  | $H_{o}$ | 0.26 | 0.27 | 0.12 | 0.31 | 0.29 | 0.01 | 0.09 | 0.02 | 0.17 | 0.13 | 0.29 | 0.26 | 0.34 | 0.52 | 0.18 | 0.33 |
|  | $H_{e}$ | 0.33 | 0.39 | 0.15 | 0.37 | 0.40 | 0.02 | 0.12 | 0.07 | 0.26 | 0.20 | 0.46 | 0.30 | 0.44 | 0.58 | 0.19 | 0.49 |
|  | HWE | *** | *** | ns | *** | *** | *** | *** | *** | *** | ns | *** | *** | *** | ns | ns | *** |
|  | $F_{IS}$ | 0.24 | 0.34 | 0.14 | 0.12 | 0.33 | 0.50 | 0.25 | 0.68 | 0.23 | 0.17 | 0.30 | 0.17 | 0.22 | 0.07 | 0.00 | 0.31 |
|  | Null | 0.07 | 0.34 | 0.03 | 0.05 | 0.10 | 0.04 | 0.06 | 0.10 | 0.19 | 0.12 | 0.40 | 0.06 | 0.09 | 0.04 | 0.00 | 0.11 |
| R2 | $N_{a}$ | 2 | 1 | 3 | 3 | 3 | 1 | 2 | 2 | 1 | 2 | 2 | 4 | 3 | 4 | 2 | 3 |
|  | $N_{e}$ | 1.03 | - | 1.32 | 2.48 | 1.02 | - | 1.01 | 1.98 | - | 1.44 | 1.13 | 2.21 | 1.13 | 1.90 | 1.01 | 2.01 |
|  | $H_{o}$ | 0.03 | - | 0.17 | 0.36 | 0.02 | - | 0.01 | 0.42 | - | 0.24 | 0.10 | 0.29 | 0.03 | 0.27 | 0.01 | 0.45 |
|  | $H_{e}$ | 0.03 | - | 0.25 | 0.60 | 0.02 | - | 0.01 | 0.50 | - | 0.31 | 0.11 | 0.55 | 0.11 | 0.48 | 0.01 | 0.50 |
|  | HWE | ns | - | *** | *** | ns | - | - | ns | - | ns | ns | *** | *** | *** | - | ns |
|  | $F_{IS}$ | -0.01 | - | 0.30 | 0.39 | -0.00 | - | - | 0.15 | - | 0.21 | 0.13 | 0.48 | 0.71 | 0.42 | - | 0.12 |
|  | Null | 0.00 | - | 0.11 | 0.17 | 0.00 | - | - | 0.05 | - | 0.26 | 0.10 | 0.18 | 0.13 | 0.14 | - | 0.03 |
| L3 | $N_{a}$ | 4 | 2 | 2 | 7 | 2 | 4 | 3 | 4 | 2 | 4 | 2 | 5 | 8 | 6 | 6 | 6 |
|  | $N_{e}$ | 1.53 | 2.00 | 1.46 | 3.72 | 1.89 | 1.29 | 2.06 | 1.25 | 1.67 | 2.83 | 1.07 | 2.42 | 4.89 | 2.68 | 2.26 | 2.97 |
|  | $H_{o}$ | 0.33 | 0.54 | 0.28 | 0.74 | 0.49 | 0.19 | 0.50 | 0.18 | 0.40 | 0.67 | 0.06 | 0.52 | 0.78 | 0.66 | 0.58 | 0.64 |
|  | $H_{e}$ | 0.35 | 0.50 | 0.32 | 0.73 | 0.47 | 0.23 | 0.51 | 0.20 | 0.40 | 0.65 | 0.06 | 0.60 | 0.80 | 0.63 | 0.56 | 0.66 |
|  | HWE | ns | ns | ns | ns | ns | *** | *** | ns | ns | ns | ns | ns | ns | ns | ns | ns |
|  | $F_{IS}$ | 0.04 | -0.07 | 0.11 | -0.00 | -0.03 | 0.16 | 0.02 | 0.10 | -0.01 | -0.03 | -0.03 | 0.11 | 0.02 | -0.05 | -0.03 | 0.03 |
|  | Null | 0.05 | 0.48 | 0.03 | 0.03 | 0.00 | 0.06 | 0.12 | 0.03 | 0.27 | 0.00 | 0.00 | 0.08 | 0.01 | 0.00 | 0.02 | 0.03 |
| L4 | $N_{a}$ | 2 | 2 | 2 | 5 | 3 | 2 | 4 | 3 | 1 | 4 | 2 | 4 | 5 | 5 | 3 | 4 |
|  | $N_{e}$ | 1.82 | 1.97 | 1.95 | 2.83 | 1.63 | 1.05 | 1.78 | 1.56 | - | 1.95 | 1.01 | 1.98 | 3.61 | 2.69 | 1.06 | 2.13 |
|  | $H_{o}$ | 0.49 | 0.39 | 0.44 | 0.62 | 0.42 | 0.04 | 0.32 | 0.38 | - | 0.51 | 0.01 | 0.55 | 0.75 | 0.49 | 0.03 | 0.50 |
|  | $H_{e}$ | 0.45 | 0.50 | 0.49 | 0.65 | 0.39 | 0.04 | 0.44 | 0.36 | - | 0.49 | 0.01 | 0.50 | 0.73 | 0.63 | 0.05 | 0.53 |
|  | HWE | ns | ns | ns | ns | ns | - | ns | ns | - | ns | - | ns | ns | ns | ns | ns |
|  | $F_{IS}$ | -0.09 | 0.20 | 0.10 | 0.04 | -0.07 | - | 0.28 | 0.05 | - | -0.04 | - | -0.10 | -0.03 | 0.23 | 0.39 | 0.06 |
|  | Null | 0.00 | 0.49 | 0.03 | 0.13 | 0.00 | - | 0.06 | 0.00 | - | 0.04 | - | 0.00 | 0.01 | 0.07 | 0.06 | 0.01 |

**Table S4**. Model-averaged survival and recapture probabilities (and their standard errors) for the three age classes (Juv, Sub and Adu) in the four SSPs (L1, L2, R1 and R2).

| Parameter | Juv | Sub | Adu |
| --- | --- | --- | --- |
| L1 |  |  |  |
| Survival | 0.35±0.06 | 0.69±0.07 | 0.70±0.02 |
| Recapture year 1 | 0.31±0.04 | 0.46±0.04 | 0.44±0.03 |
| Recapture year 2 | 0.39±0.04 | 0.55±0.04 | 0.53±0.02 |
| Recapture year 3 | 0.29±0.04 | 0.44±0.05 | 0.42±0.03 |
| Recapture year 4 | 0.13±0.03 | 0.23±0.04 | 0.21±0.03 |
| Recapture year 5 | 0.25±0.07 | 0.39±0.09 | 0.37±0.08 |
| Recapture year 6 | 0.32±0.05 | 0.48±0.05 | 0.46±0.04 |
| Recapture year 7 | 0.25±0.04 | 0.40±0.04 | 0.38±0.03 |
| Recapture year 8 | 0.32±0.04 | 0.48±0.05 | 0.46±0.03 |
| Recapture year 9 | 0.31±0.04 | 0.46±0.04 | 0.44±0.03 |
| L2 |  |  |  |
| Survival | 0.46±0.01 | 0.58±0.01 | 0.60±0.01 |
| Recapture year 1 | 0.41±0.03 | 0.29±0.03 | 0.32±0.01 |
| Recapture year 2 | 0.27±0.01 | 0.20±0.01 | 0.25±0.01 |
| Recapture year 3 | 0.37±0.02 | 0.27±0.01 | 0.27±0.01 |
| Recapture year 4 | 0.19±0.02 | 0.20±0.01 | 0.20±0.01 |
| Recapture year 5 | 0.14±0.05 | 0.31±0.02 | 0.34±0.01 |
| R1 |  |  |  |
| Survival | 0.69±0.03 | 0.87±0.02 | 0.82±0.01 |
| Recapture year 1 | 0.76±0.02 | 0.77±0.02 | 0.74±0.01 |
| Recapture year 2 | 0.75±0.03 | 0.77±0.02 | 0.74±0.01 |
| Recapture year 3 | 0.63±0.03 | 0.65±0.03 | 0.61±0.02 |
| Recapture year 4 | 0.81±0.02 | 0.82±0.02 | 0.79±0.02 |
| Recapture year 5 | 0.79±0.03 | 0.80±0.03 | 0.77±0.02 |
| R2 |  |  |  |
| Survival | 0.64±0.02 | 0.76±0.02 | 0.78±0.01 |
| Recapture year 1 | 0.61±0.02 | 0.58±0.02 | 0.68±0.01 |
| Recapture year 2 | 0.63±0.02 | 0.59±0.02 | 0.69±0.01 |
| Recapture year 3 | 0.67±0.02 | 0.64±0.02 | 0.73±0.01 |
| Recapture year 4 | 0.65±0.03 | 0.62±0.02 | 0.72±0.01 |
| Recapture year 5 | 0.67±0.03 | 0.63±0.03 | 0.73±0.02 |
